# Multiple routes of adaptation to high levels of CIN and aneuploidy in budding yeast

**DOI:** 10.1101/2022.04.21.489003

**Authors:** Matthew N. Clarke, Theodor Marsoner, Manuel Alonso Y Adell, Madhwesh C. Ravichandran, Christopher S. Campbell

## Abstract

Both an increased frequency of chromosome missegregation (chromosomal instability) and the presence of an abnormal complement of chromosomes (aneuploidy) are hallmarks of cancer. Paradoxically, both chromosomal instability and aneuploidy are also associated with substantial decreases in cellular fitness. To better understand how cells are able to adapt to high levels of chromosomal instability, we previously examined yeast cells that were deleted of the gene *BIR1*, a member of the chromosomal passenger complex (CPC). The CPC is an essential regulator of chromosome segregation fidelity. We found *bir1Δ* cells quickly adapted by acquiring specific combinations of beneficial aneuploidies. However, targeted mutations of specific genes were notably absent in the short term. In this study, we monitored these yeast strains for longer periods of time to determine how cells adapt to high levels of both CIN and aneuploidy in the long term. We identify suppressor mutations that mitigate the chromosome missegregation phenotype. The mutated proteins fall into four main categories: outer kinetochore subunits, members of the SCF^Cdc4^ complex, the mitotic kinase Mps1, and a member of the CPC itself. These mutants function in two distinct ways, as mutations in the outer kinetochore suppress Bir1 deletion indirectly by destabilizing connections between the chromosomes and the mitotic spindle, whereas the other three categories of mutations affect the CPC directly. As a consequence of the accumulation of suppressor point mutations, overall levels of aneuploidy decreased. These experiments demonstrate a timeline of adaptation to high rates of CIN wherein cells first acquire specific aneuploidies that suppress the CIN phenotype, next develop point mutations that more specifically target the source of CIN, and finally reduce the level of aneuploidy to relieve the fitness burden placed by aneuploidy on the cell.

## Introduction

The accurate distribution of chromosomes to daughter cells is a fundamental requirement of cell division. An increase in the frequency of errors in chromosome segregation is called chromosomal instability (CIN). CIN leads to abnormal karyotypes through the gain or loss of chromosomes, a state called aneuploidy. Aneuploidy and CIN are both hallmarks of cancer that have causative roles in cancer development, cancer progression, and resistance to chemotherapy (reviewed in Ben-David and Amon, 2020). Despite the promotion of cell proliferation in cancer, aneuploidy and CIN have consistently been demonstrated to decrease cell growth and division (Gropp et al., 1983; Torres et al., 2007). Cancers therefore likely develop adaptations to ameliorate the negative affects of CIN and aneuploidy.

Different models for how cells adapt to CIN and aneuploidy have been proposed. Cells could adapt via compensatory mutations that decrease the levels of CIN after the accumulation of beneficial aneuploidies (Cahill et al., 1999; Sansregret et al., 2017; Kwon et al., 2008). Alternatively, it has been postulated that cancer cells could adapt to CIN and aneuploidy through mutations that lead to aneuploidy tolerance (Torres et al., 2010). Additionally, it has been suggested that CIN and aneuploidy provide for a fast, but transient mechanism of adaptation. In this model, aneuploidy provides a short term benefit that outweighs its downsides, but would eventually be superceded by more targeted genome alterations (Yona et al., 2012). How aneuploidy and more specific types of mutations affect each other during the course of adaptation is currently unknown.

The molecular mechanisms that lead to CIN in cancer cells have long been elusive (Gordon et al., 2012). However, one relevant phenotype that is frequently observed across many cancer types is the overstabilization of connections between microtubules and kinetochores, which are the binding sites for microtubules at the centromeres of chromosomes. These overstabilized attachments lead to the accumulation of misattached chromosomes where both of the sister chromatids are attached to microtubules emanating from the same spindle pole (merotelic and syntelic attachments) (Bakhoum et al., 2009a). One or both of the kinetochores must then be detached from the microtubules in order to properly distribute one sister chromatid to each daughter cell. Although the basis behind this phenotype in cancer cells is currently unknown, the central player in destabilizing erroneous kinetochore-microtubule attachments is the chromosomal passenger complex (CPC). The CPC contains a kinase, Aurora B, that phosphorylates kinetochores to lower their affinity for microtubules (Tanaka et al., 2002; Cheeseman et al., 2006; Sarangapani et al., 2013; Kalantzaki et al., 2015). Inhibition of Aurora B in mammalian cells leads to an increased frequency of lagging chromosomes in anaphase due to the inability to detach microtubules from kinetochores that are attached to both spindle poles (Cimini et al., 2006). This lagging chromosome phenotype is also frequently observed in cancers (Bakhoum et al., 2014).

In addition to Aurora B, the CPC contains the subunits INCENP, Survivin, and Borealin. The C-terminus of INCENP binds to and activates Aurora B, while the N-terminus binds to Survivin and Borealin (Bishop and Schumacher, 2002; Honda et al., 2003; Sessa et al., 2005; Ainsztein et al., 1998; Klein et al., 2006; Jeyaprakash et al., 2007). Survivin and Borealin target the complex to centromere-proximal chromatin through an interaction with Shugoshin. This interaction has been shown to promote the activity of the CPC in correcting erroneous kinetochore-microtubule attachments through phosphorylation of substrates at the outer kinetochore (Gassmann et al., 2004; Vanoosthuyse et al., 2007; Kawashima et al., 2010; 2007).

The budding yeast *S. cerevisiae* is a valuable model organism for studying CIN and aneuploidy as it can tolerate high levels of CIN that are similar to what is frequently observed in cancer. We previously determined how cells initially adapt to extremely high rates of chromosome missegregation by growing budding yeast cells carrying mutations that decrease CPC function (Ravichandran et al., 2018). The budding yeast homologs of the CPC subunits Aurora B, INCENP, Survivin and Borealin are Ipl1, Sli15, Bir1, and Nbl1, respectively (Figure 1A). By sequencing populations of yeast that were grown in the absence of the CPC subunit Survivin/Bir1, we found that the yeast adapted by acquiring specific aneuploidies that decreased the rate of CIN (Ravichandran et al., 2018). The compositions of karyotypes were further refined over time until the cells acquired certain combinations of beneficial aneuploid chromosomes. However, these adapted cells were still substantially less fit than wild-type due to a combination of the negative affects of the acquired aneuploidies and residual chromosomal instability. It is currently not known how cells with high rates of CIN and aneuploidy further adapt after the initial optimization of their karyotypes.

**Figure 1.**
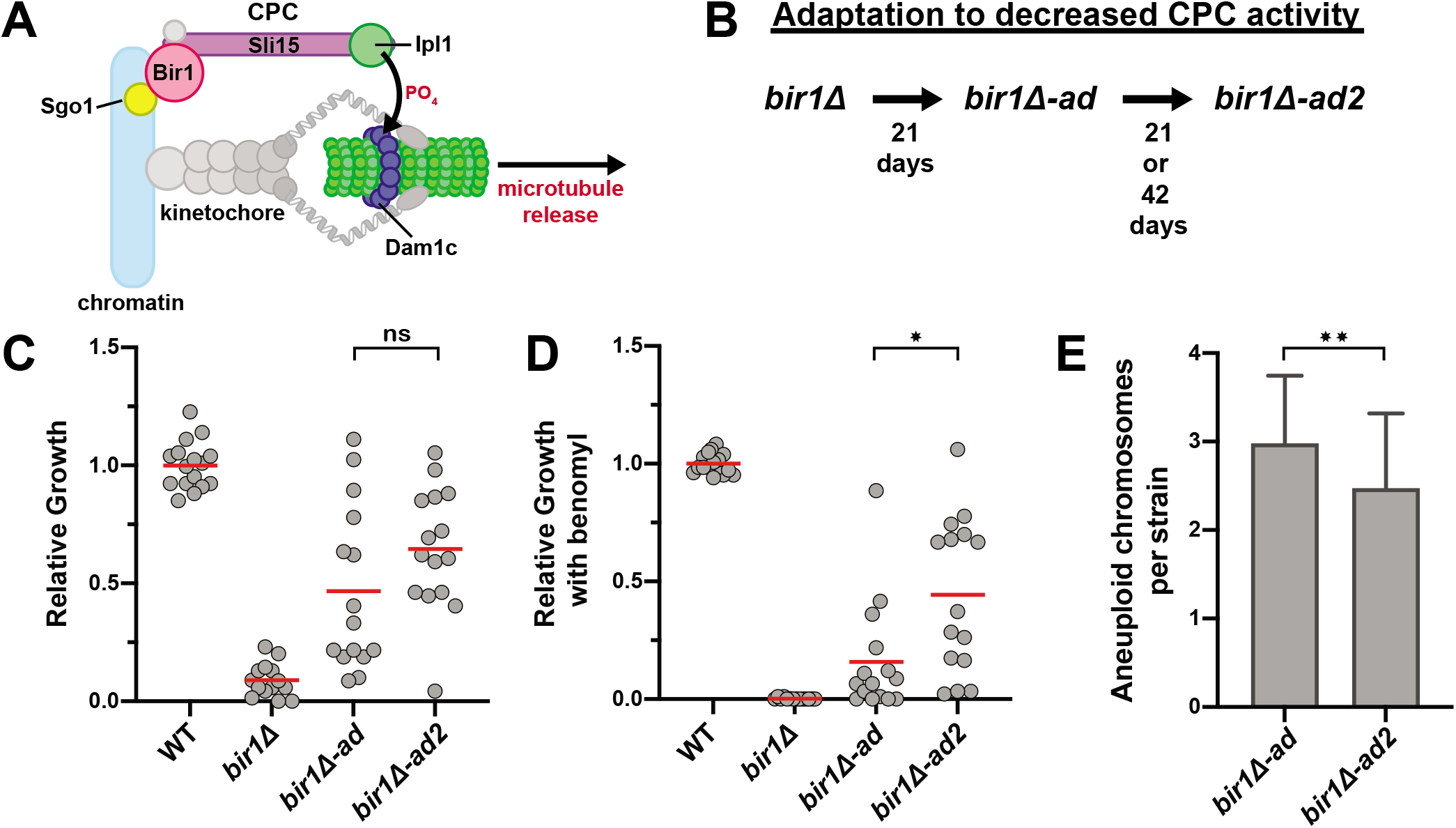
Strains with high levels of CIN and aneuploidy adapt by decreasing CIN. (A) Schematic summarizing the main localization of the CPC and its role in phosphorylating the Dam1 complex (Dam1c) to destabilize microtubule attachments. (**B**) Summary of the adaptation timeline. *bir1*Δ strains from tetrads were grown clonally for 21 days to produce *bir1*Δ-*ad* strains. Colonies from the *bir1*Δ-*ad* strains were then grown in liquid media for 21 or 42 additional days to produce the *bir1*Δ-*ad2* strains. (**C** and **D**) Growth comparisons of wild-type (WT), unadapted (*bir1*Δ), partially adapted (*bir1*Δ-*ad*), and further adapted (*bir1*Δ-*ad2*) strains as measured by area of colony growth after serial dilution. 10-fold serial dilutions were made on YPAD plates containing either 0.1% DMSO (**C**) or 10μg/mL of benomyl (**D**). The mean (red line) is shown. (**E**) Comparison of the total number of aneuploid chromosomes per strain for the *bir1*Δ-*ad* (47 strains) and *bir1*Δ-*ad2* (53 strains) collections. Only data from *bir1*Δ-*ad* strains that were subjected to additional adaptation are shown. (ns) not significant; (*) P<0.05; (**) P<0.01; unpaired t-test.

In this study, we have now monitored these CPC deficient cells for longer periods of time to determine how cells evolve to cope in the long term with increased levels of both CIN and aneuploidy. We aimed to determine if they adapt through genetic changes that either i) allow for better aneuploidy tolerance, ii) decrease the levels of CIN, or iii) further optimize their karyotypes. We found that cells evolved through hypomorphic mutations in essential genes that decreased the levels of CIN. The lower levels of CIN then allowed the cells to decrease their levels of aneuploidy, as the beneficial affects of the extra chromosomes no longer outweighed the fitness costs. The identified mutations fall into two broad functional categories. The first category of mutations destabilizes kinetochore-microtubule interactions, counteracting the overstabilization created by decreased CPC activity. These mutations were found in outer kinetochore proteins that directly interact with microtubules. The second category of mutations affect the function of the CPC more directly. These include mutations in the CPC itself, the mitotic kinase Mps1, and the ubiquitin ligase SCF^Cdc4^ complex. SCF^Cdc4^ mutations increase the recruitment of the CPC to key regulatory regions in both yeast and human cells. We conclude that cells generally adapt to high levels of CIN and aneuploidy through mutations that alleviate the CIN phenotype. Further, we have identified specific pathways of adaptation for defects in CPC function.

## Results

### Adaptation to high levels of CIN and aneuploidy occurs through point mutations that reduce the rate of CIN

To determine how cells adapt to high rates of CIN and aneuploidy, we started with a collection of haploid yeast strains that had previously been cultured for 21 days via clonal expansion following Bir1 deletion (Ravichandran et al., 2018). We call these partially adapted strains *bir1Δ-ad*. These strains have a high frequency of specific aneuploidies that decrease the rate of CIN (Figure S1A). However, they are not enriched for mutations that are related to CIN (Figure S1B). We adapted 68 of these strains for an additional 21 or 42 days (*bir1Δ-ad2*) in liquid culture to determine how cells continue to evolve after their initial adaptation through aneuploidy (Figure 1B). In comparison to the clonal expansion of the initial adaptation, liquid culture adaptation allows time for rarely occurring beneficial genomic changes to take over the population competitively. This additional adaptation greatly improved the growth of some strains, but did not result in a significant level of growth improvement for all of the adapted populations combined (Figure 1C). Cells with impaired chromosome segregation are especially sensitive to microtubule depolymerizing drugs. Consistent with this, Bir1 deletion results in a strong sensitivity to moderate amounts of the microtubule-depolymerizing drug benomyl (Makrantoni and Stark, 2009). Many of the *bir1Δ-ad2* strains have strongly decreased benomyl sensitivity in comparison to the *bir1Δ-ad strains*, suggesting that they have acquired additional changes that suppress the CIN phenotype (Figure 1D). To determine if this additional adaptation occurs through further increased aneuploidy, we measured chromosome copy numbers via read counts from whole genome sequencing. Overall, the *bir1Δ-ad2* strains have decreased numbers of aneuploid chromosomes, suggesting that these strains do not adapt through further increases in aneuploid chromosomes that attenuate CIN (Figures 1E and S1A). We also did not observe any partial-chromosome copy number changes in the adapted strains. We conclude that the additional adaptation to CIN in the further adapted CPC-deficient strains likely comes from specific mutations rather than chromosome copy number alterations.

Adaptation to CIN and aneuploidy could potentially come from mutations that either decrease CIN or increase aneuploidy tolerance. To determine if mutations in the adapted strains fall into these categories, we searched for non-synonymous mutations that arose during liquid culture adaptation in the whole genome sequencing data. We identified 97 such mutations in 68 strains. These mutations were enriched in genes related to CIN (Figure S1B). Mutations that increase aneuploidy tolerance have previously been reported in budding yeast (Torres et al., 2010). However, there was no overlap between the 22 genes identified in the aneuploidy tolerance screen and the genes mutated in the *bir1Δ-ad2* strains. We therefore used the most heavily characterized gene whose loss leads to aneuploidy tolerance, the ubiquitin-specific protease Ubp6 (Torres et al., 2010). We tested if a mutation in Ubp6 that was previously reported to increase aneuploidy tolerance would increase resistance to *bir1Δ*. The mutation was introduced into a haploid strain whose only copy of *BIR1* is on a minichromosome that also contains the *URA3* gene. This minichromosome can be selected against by using 5-Fluoroorotic acid (5-FOA), which is converted into a toxin by the *URA3* gene product. Addition of the *ubp6(E256X)* mutation decreased, rather than increased, viability after Bir1 was deleted (Figure S1C). This result suggests that aneuploidy tolerance does not lead to CIN tolerance following Bir1 deletion. We conclude that cells with high levels of CIN and aneuploidy adapt primarily through mutations that mitigate CIN rather than aneuploidy.

### *bir1Δ* supressor mutations fall into four major categories

To determine which categories of mutations are most prevalent in our adapted strains, we searched for the enrichment of functional gene ontology (GO) terms. We found significant enrichment of genes related to “chromosome segregation” (FDR=3.53×10^-5^) and “SCF-dependent proteasomal ubiquitin-dependent protein catabolic process” (FDR=1.3×10^-1^, Table S2). Further refinement of these categories revealed that the mutations fall largely into four categories (Table 1).

**Table 1.** Candidate mutations identified in *bir1*Δ-*ad2*.

| Gene mutated | Residue changes | Tested | Chromosome | Days of liquid adaptation | Essential | Function |
| --- | --- | --- | --- | --- | --- | --- |
| *DUO1* | P17L | Yes | 7 | 21 | Yes | Kinetochore-microtubule attachment |
| *DAD1* | N43S | Yes | 4 | 21 | Yes |  |
| *DAD2* | K11Q | Yes | 11 | 21 | Yes |  |
| *ASK1* | S216F | Yes | 11 | 21 | Yes |  |
| *SPC34* | D119A | Yes | 11 | 21 | Yes |  |
| *NDC80* | K181N | Yes | 9 | 42 | Yes |  |
| *SPC105* | R583G<sup>b</sup> | Yes | 7 | 21 | Yes |  |
| *SLI15* | L71S | Yes | 2<sup>a</sup> | 42 | Yes | CPC subunit |
| *SLI15* | P109A | No | 2<sup>a</sup> | 21 | Yes |  |
| *SLI15* | G334S | Yes | 2<sup>a</sup> | 21 | Yes |  |
| *CDC53* | K448E | No | 4 | 21 | Yes | S-phase entry |
| *CDC53* | A486P | No | 4 | 21 | Yes |  |
| *CDC34* | M64T<sup>b</sup> | Yes | 4 | 21 | Yes |  |
| *CDC4* | S438G | Yes | 6 | 21 | Yes |  |
| *CDC4* | G439S | Yes | 6 | 21 | Yes |  |
| *CDC4* | G652D | No | 6 | 21 | Yes |  |
| *MPS1* | R596H | Yes | 4 | 21 | Yes | Initiation of the SAC |
| *MPS1* | W629S | No | 4 | 21 | Yes |  |
| *MPS1* | V631M | Yes | 4 | 21 | Yes |  |
| *MIF2* | D241N | Yes | 11 | 42 | Yes | Inner kinetochore |
| *RTG2* | G154S | Yes | 7 | 21 | Yes | Mitochondrial sensor |
| *RTG2* | H425L | No | 7 | 21 | Yes |  |
| *RTG2* | A433P | No | 7 | 21 | Yes |  |
| *SPC97* | S816X | Yes | 8<sup>a</sup> | 21 | Yes | Spindle pole body |

<sup>a</sup> chromosomes that are commonly aneuploid in *bir1Δ* strains
<sup>b</sup> identified in the same strain

The first category is in proteins that form the outer kinetochore, including mutations in the Dam1 complex, the Ndc80 complex, and Spc105/KNL1. All three of these proteins/protein complexes directly bind to microtubules. The Dam1 complex was the most heavily represented in the data set with mutations in 5 of the 10 subunits. Mutations in the Dam1 complex subunit Dam1 that mimic Ipl1 phosphorylation have been previously shown to suppress CPC mutations (Cheeseman et al., 2002). However, none of the mutations identified in the *bir1Δ-ad2* strains were in the Dam1 protein itself, indicating that they do not directly mimic similar phosphorylation events (Figure 2A).

**Figure 2.**
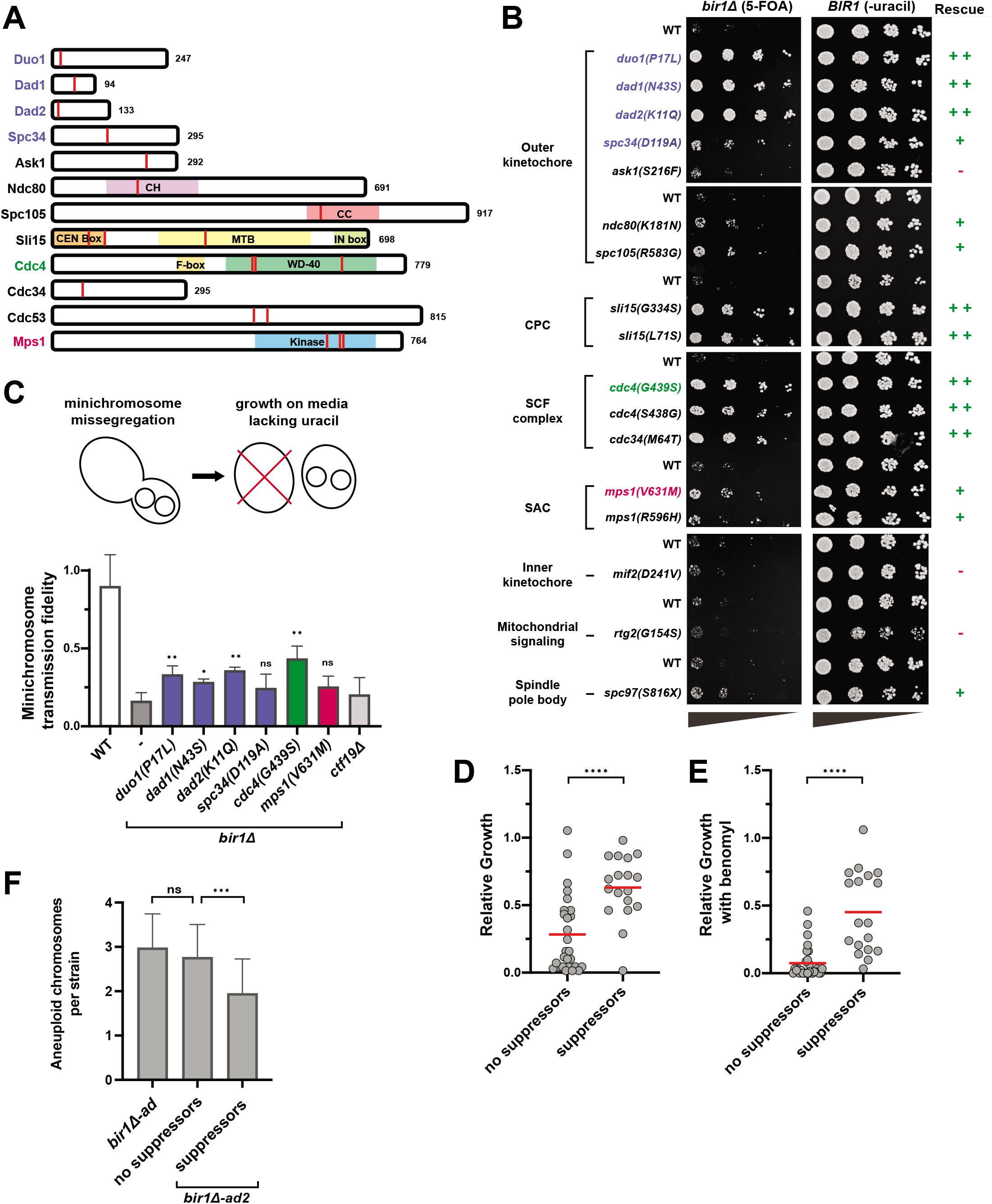
Four main categories of mutations rescue Bir1 deletion. (**A**) Schematics showing the location of mutations from Table 1 (red lines). Domains of interest were identified using the Saccharomyces Genome Database. (**B**) Serial dilutions of strains engineered with the indicated mutations identified in *bir1*Δ-*ad2* strains were tested for rescue of Bir1 deletion. 10-fold serial dilutions on the indicated media are shown. The mutations colored in blue, green, or red were selected for additional characterization. (**C**) Top: schematic summarizing how loss of the *URA3*-containing plasmid is used to measure relative missegregation rates. Bottom: Proportion of colonies that grow on plates with restrictive (lacking uracil) versus permissive (YPAD) media. Many of the suppressor mutants significantly increase the segregation fidelity of the minichromosome in a *bir1*Δ background. *ctf19*Δ serves as a positive-control for decreased minichromosome transmission fidelity. Statistical significance is relative to *bir1*Δ alone. (**D** and **E**) Growth comparisons of *bir1*Δ-*ad2* strains that contain (18 strains) or do not contain (28 strains) confirmed suppressor mutations as measured by area of colony growth after serial dilution. 10-fold serial dilutions were made on YPAD plates containing either 0.1% DMSO (**D**) or 10μg/mL of benomyl (**E**). (**F**) The strains with confirmed suppressors had significantly reduced levels of aneuploidy. (ns) not significant; (*) P<0.05; (**) P<0.01; (***) P<0.001; (****) P<0.0001; unpaired t-test.

The second category of mutations is the CPC subunit Sli15. Mutations in Sli15 that disrupt binding to Bir1 or prevent Cdc28 (cyclin-dependent kinase) phosphorylation have both been previously shown to suppress Bir1 deletion (Campbell and Desai, 2013). All three of the observed mutations are consistent with these previous observations, as two were in the CEN box that interacts with Bir1 (L71S and P109H) and one was directly adjacent to the main Cdc28 phosphorylation site (G334S).

The third category of common mutations in the adapted strains is in subunits of the SCF ubiquitin-protein ligase complex. The SCF is primarily involved in the initiation of S-phase, but it also has roles in mitosis (Goh and Surana, 1999). We identified mutations in the core SCF subunits Cdc53 and Cdc34 as well as the F-box adaptor protein Cdc4. F-box proteins dictate the substrate specificity of the SCF complex (Jonkers and Rep, 2009). Important substrates of SCF^Cdc4^ include the cell cycle kinase inhibitors Sic1 and Far1. No mutations in any other F-box proteins were identified, suggesting that the ability to suppress Bir1 deletion may be specific to SCF^Cdc4^. All three Cdc4 mutations were in the WD-40 domain, which directly interacts with SCF complex substrates (Figure 2A). No direct connection between the SCF complex and the CPC has been previously identified.

The final category of potential suppressor mutations was in the mitotic kinase Mps1. All three of the mutations in Mps1 were in the kinase domain (Figure 2A). Mps1 has functions in spindle assembly checkpoint signaling, correction of misattached chromosomes, and spindle pole body duplication (reviewed in Liu and Winey, 2012). We were surprised to identify mutations in Mps1 as it would be expected to promote rather than antagonize CPC activity, and mutations in Mps1 that suppress CPC deficiency have not previously been identified.

Intriguingly, all of the mutations in Table 1 are in essential genes, so they are most likely to be hypomorphic alleles. To determine if the identified mutations are sufficient to rescue *bir1Δ*, we introduced mutations from each category prior to Bir1 deletion. In addition, we included mutations identified in the adapted strains in three other potentially interesting proteins: the inner kinetochore protein Mif2, the spindle pole body protein Spc97, and the mitochondrial signaling protein Rtg2. Rtg2 was selected because we identified three independent mutations in this protein. After selection of *bir1Δ* on 5-FOA plates, all of the tested mutations at least partially rescued growth with the exception of *mif2, rtg2*, and *ask1* (Figure 2B). We identified multiple mutations that suppress the Bir1 deletion growth phenotype for each of the four major categories. To determine if the mutants were able to rescue the chromosome missegregation phenotype of Bir1 deletion, we measured the fidelity of minichromosome transmission with representative suppressor mutations from the outer kinetochore, SCF complex, and Mps1 categories. We did not perform any further experiments with the Sli15 mutations, as similar mutations have previously been characterized (Campbell and Desai, 2013). For the SCF complex, we analyzed mutations that we identified in Cdc4, as mutations in the F-box subunit are less likely to have pleiotropic effects. Minichromosome transmission fidelity was higher than the *bir1Δ* control for mutations in all three categories, although the degree of rescue for the Mps1 mutation was not significant (p=0.1, Figure 2C). We have therefore identified four categories of mutations – in the outer kinetochore, the CPC, the SCF, and Mps1 – that are frequently mutated to suppress the chromosome missegregation phenotype of yeast with impaired CPC activity.

We next wanted to determine the degree to which the identified mutations contribute to adaptation. We compared the growth of *bir1Δ-ad2* strains that either do or do not contain identified suppressor mutations. All mutations listed in Table 1 except for those in *MIF2*, *RTG2*, and *ASK1* were considered suppressors for this analysis. On average, adapted strains that contain identified suppressor mutations grew substantially better than those without suppressors, suggesting that we identified most of the impactful mutations and that they contributed greatly to the adaptation (Figures 2D and 2E). This difference was significant either with or without the addition of benomyl. Furthermore, the degree of aneuploidy is significantly reduced in the strains with suppressor mutations, indicating that these mutations decreased the need for aneuploid chromosomes to suppress the *bir1Δ* phenotype (Figure 2F). This decrease was seen across all of the observed aneuploid chromosomes (Figure S1D). The aneuploidy burden in the further adapted strains was therefore reduced by decreasing the requirement for aneuploidy rather than decreasing the impact of aneuploidy on cellular fitness. These results provide a timeline of events for adaptation to high rates of CIN. First, the cells acquire specific aneuploidies that suppress the CIN phenotype. The cells then acquire optimal combinations of aneuploidies (Ravichandran et al., 2018). Next, point mutations arise that more specifically target the source of the CIN leading to a reduction of CIN. Finally, the level of aneuploidy decreases to relieve the fitness burden placed on the cell.

### Suppressor mutations in the Dam1 complex create unattached kinetochores

We next sought to determine where in the chromosome biorientation pathway each of these categories of mutations falls. Defects in CPC activity lead to the overstabilization of kinetochore-microtubule attachments and the failure to correct attachments where both sister chromatids attach to microtubules emanating from the same pole (syntelic attachments) (Biggins et al., 2001; Pinsky et al., 2006). Mutations that rescue CPC defects are therefore likely to restore the higher turnover of kinetochore-microtubule attachments. This could occur either by increasing the activity of the CPC or by decreasing the microtubule binding activity of kinetochores. To determine if the mutations act downstream of the CPC, we tested if they could rescue a temperature-sensitive mutation in the CPC kinase Ipl1/Aurora B. Of the mutations tested, only the Dam1c mutants significantly rescued *ipl1-321* at the restrictive temperature (Figure 3A). This result indicates that the Dam1c mutations affect the pathway downstream of the CPC, whereas the SCF complex and Mps1 potentially affect the CPC itself.

**Figure 3.**
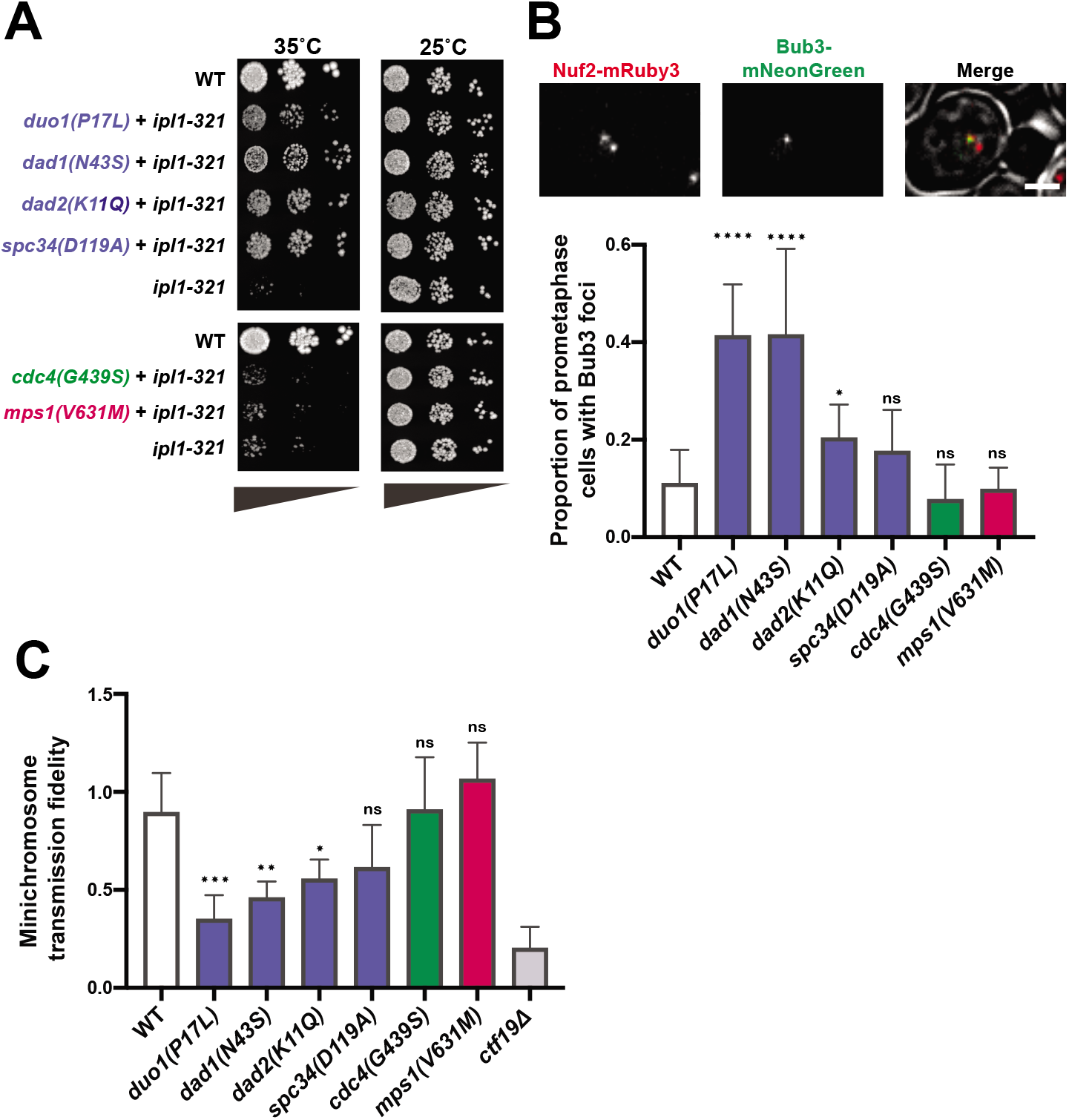
Suppressor mutations in the Dam1 complex result in elevated spindle assembly checkpoint activity and minichromosome missegregation. (**A**) Serial dilutions on YPAD media at the permissive (25°C) or restrictive (35°C) temperatures for the *ipl1-321* mutation. The suppressor mutations from the Dam1c (blue) were the only mutants that rescued viability at the restrictive temperature. (**B**) Top: representative images showing Bub3-mNeonGreen localization to kinetochores (Nuf2-mRuby3), indicating spindle assembly checkpoint activation in wild-type yeast. Bottom: quantification of the proportion of prometaphase cells with Bub3-mNeonGreen foci. Only the mutations in Dam1 complex (blue) show a significant increase in SAC activity. (**C**) Proportion of colonies that grow on plates with restrictive (lacking uracil) versus permissive (YPAD) media after 48 hours growth under permissive conditions with a *URA3*-containing plasmid. The suppressor mutations in the Dam1c (blue) show a significant reduction of minichromosome segregation fidelity. (ns) non-signifcant; (*) P<0.05; (**) P<0.01; (***) P<0.001; (****) P<0.0001; unpaired t-test.

If the mutations rescue by globally destabilizing kinetochore-microtubule attachments, then the number of unattached kinetochores should increase even in the presence of Bir1. Since unattached kinetochores trigger the spindle assembly checkpoint, we first determined if the mutations induce a delay in mitosis. None of the identified mutations showed substantial changes in cell cycle duration, as measured by the sustained accumulation of large budded cells over time after release from synchronization in G1 (Figure S2A). These mutations therefore do not maintain sustained checkpoint activity. As a more sensitive assay, we monitored the presence of unattached kinetochores by measuring the frequency of cells with foci of the checkpoint protein Bub3 that colocalize with kinetochore clusters. For this assay, we only counted cells that have two kinetochore clusters that are separated yet still in close proximity, indicative of a stage around the prometaphase to metaphase transition. At this stage, three of the four Dam1c mutations resulted in significantly more cells with Bub3-mNeonGreen foci than the wild-type control (Figure 3B). By contrast, suppressor mutations in Cdc4 and Mps1 did not increase the frequency of Bub3 foci. We conclude that suppressor mutations in the Dam1 complex transiently increase the number of unattached kinetochores, likely through increased kinetochore-microtubule attachment turnover.

The suppression of the *bir1Δ* chromosome missegregation phenotype through an increase in unattached kinetochores in Dam1 complex mutants suggests that there is a restoration of the balance between microtubule attachment and detachment when both mutations are combined. If this were the mechanism of rescue, then we would expect that in the absence of Bir1 deletion, the increase in kinetochore-microtubule attachment turnover in the Dam1 mutants would overly destabilize attachments and increase the chromosome missegregation rate. To test this, we measured the rate of minichromosome transmission in strains that have suppressor mutations and wild-type *BIR1*. Similar to the results of the Bub3 foci counts, three of the four Dam1c mutants showed a significant decrease in minichromosome transmission fidelity (Figure 3C). The Cdc4 and Mps1 mutations had no measurable effect. The Dam1c mutation that did not show a significant result in either the unattached kinetochore or chromosome segregation assays, *spc34*(G439S), also had the weakest rescue of growth following Bir1 deletion (Figure 2B). Overall, these results demonstrate that mutations in outer kinetochore proteins rescue deficient CPC activity by destabilizing kinetochore-microtubule attachments. In contrast, Cdc4 and Mps1 suppressor mutations act through an alternative mechanism that does not directly affect the stability of kinetochore-microtubule attachments.

### Dam1c suppressor mutations have similar phenotypes to a phosphomimic mutation that decreases Dam1c microtubule binding

Phosphomimics of Ipl1 phosphorylation sites on the Dam1 subunit of the Dam1 complex have previously been shown to partially rescue *ipl1-ts* mutants (Cheeseman et al., 2002). These phosphosites include serine 20 (*dam1(S20D)*) near the N-terminus and three serines (S257, S265, S292, *dam1(3D)*) closer to the C-terminus of the protein (Figure 4A). The two phosphomimic mutants change the serines to negatively charged aspartic acid residues. To determine if the identified suppressor mutations function similarly to the phosphomimics, we first determined if the *dam1(S20D)* or *dam1(3D)* mutations rescue Bir1 deletion. Intriguingly, *dam1(S20D)* rescues Bir1 deletion but *dam1(3D)* does not, indicating that these mutants are mechanistically distinct (Figure 4B). Both mutations increase the frequency of Bub3 foci in prometaphase cells, similarly to the Dam1c suppressor mutations that we identified (Figure 4C). However, the *3D* mutant shows a strong cell cycle delay, further indicating that it differs from the other Dam1c mutants in either mechanism or severity (Figure 4D). Surprisingly, despite the delay, the *3D* mutant does not affect minichromosome transmission fidelity (Figure 4E). The *dam1*(*S20D*) mutation, however, does show a decrease in minichromosome transmission in line with the suppressor mutations (Figure 4E). We conclude that the mutations in the Dam1 complex that adapt to suppress Bir1 deletion have a similar phenotype to the *dam1(S20D)* mutation. Intriguingly, the *S20D* phosphomimic mutation has been shown to disrupt microtubule binding of kinetochores *in vitro*, whereas the triple serine phosphomimic (*3D*) mutant maintained wild-type microtubule binding activity (Sarangapani et al., 2013). The similarity in phenotype between the suppressor mutations and *dam1(S20D)* suggests that the suppressor mutations may also directly decrease kinetochore-microtubule affinity.

**Figure 4.**
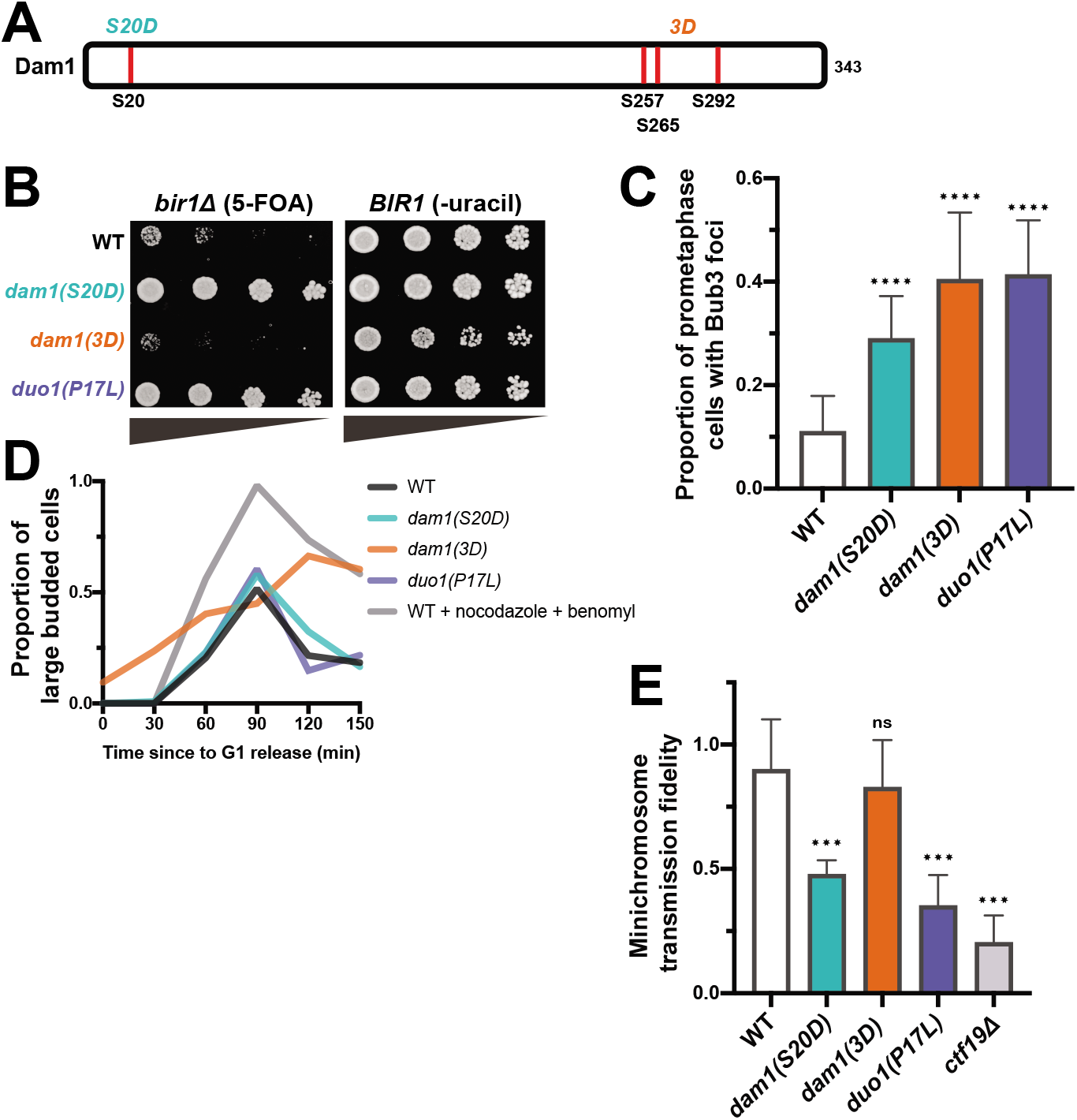
Suppressor mutations in the Dam1 complex have phenotypes similar to the *dam1(S20D)* phosphomimic mutation. (**A**) A schematic showing the relative locations of the Ipl1 phosphosites in Dam1 that are mutated to aspartic acid in the phosphomimics. (**B**) Serial dilutions of strains engineered with the indicated mutations were tested for rescue of Bir1 deletion. 10-fold serial dilutions on the indicated media are shown. *dam1(S20D)* rescues loss of CPC activity to a similar extent as a suppressor mutant in Duo1. (**C**) Quantification of the proportion of prometaphase cells with Bub3-mNeonGreen foci localized to kinetochores (Nuf2-mRuby3), indicating SAC activation in all three mutant strains. (**D**) Degree of cell cycle delay as measured by percentage of large budded cells over time after release from G1 arrest. Wild-type (WT) cells treated with nocodazole and benomyl were used as a positive control for mitotic arrest. (**E**) Proportion of colonies that grow on plates with restrictive (lacking uracil) versus permissive (YPAD) media after 48 hours growth under permissive conditions with a *URA3*-containing plasmid. Minichromosome segregation fidelity is reduced in the *dam1(S20D)* strain, similar to the *duo1(P17L)* strain. (ns) non-signifcant; (***) P<0.001; (****) P<0.0001; unpaired t-test.

If both the Dam1c suppressor mutations and *dam1(S20D)* act through destabilizing kinetochore-microtubule attachments, we hypothesized that combining the two mutants might destabilize the connections to a greater extent. Indeed, combination of either *duo1(P17L)* or *dad2(K11Q)* with *dam1(S20D)* resulted in zero viable spores after three days if growth (Figure S2B). For the *duo1(P17L)*, *dam1(S20D)* double mutant, rare colonies grew up after five days. These double mutant cells displayed a strong metaphase delay and severe minichromosome loss, as expected for high rates of unattached kinetochores (Figures S2C and S2D). We conclude that the suppressor mutations in the Dam1 complex and the *dam1(S20D)* mutant both act by destabilizing kinetochore-microtubule attachments. The combination of the mutants increases the severity of the phenotype to create an extended spindle assembly checkpoint arrest and cell death.

### Dam1c suppressor mutations lie proximal to multimerization interfaces and decrease spindle localization in vivo

To determine a potential mechanism of action for the Dam1c suppressor mutations, we mapped them onto the existing crystal structure of the *C. thermophilum* version of the complex (Jenni and Harrison, 2018). Three of the mutations are in helices that lie near interfaces of oligomerization between Dam1c decamers. *spc34(D119A)* is found in a region that corresponds to a short helix that forms part of interface 1. The *dad1(N43S)* mutation affects a highly conserved residue that lies on the surface of interface 2 (Figure 5A). The *dad2(K11Q)* mutation affects a highly conserved lysine that is also proximal to interface 2, but is not directly on the surface. The *duo1(P17L)* mutation lies in a region outside of the crystal structure. However, the affected residue would likely be near the N-terminal part of Duo1, which resides in interface 1. These interfaces are proposed to be important for the ring formation that allows the complex to encircle microtubules. It has been noted that serine 20 of the Dam1 protein is potentially also located in the vicinity of interface 1 (Jenni and Harrison, 2018). These results suggest that the suppressor mutants could act by limiting higher-order oligomerization of the Dam1 complex.

**Figure 5.**
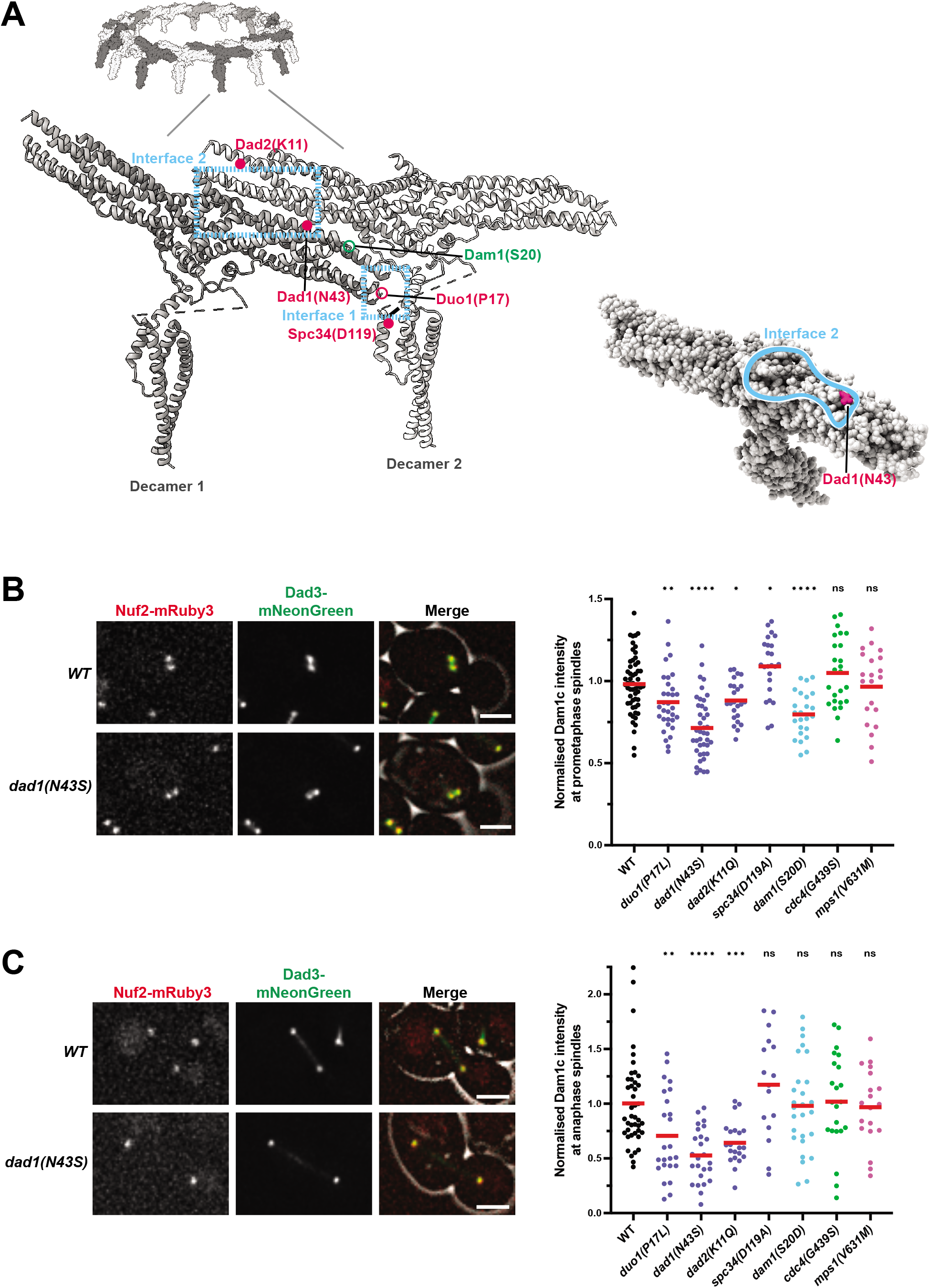
Suppressor mutations in the Dam1 complex reduce spindle localization. (**A**) View of the *C. thermophilum* Dam1c-ring, with focus on two adjacent protomers (labeled decamers 1 and 2). Interfaces 1 and 2 between adjacent decamers are shown with the blue dashed boxes as previously identified by Jenni and Harrison (Jenni and Harrison, 2018). Mutations in the *S. cerevisiae* proteins have been labeled based on sequence homology with *C. thermophilum*. Mutations within the structure are depicted as solid circles. Those that are located outside of the structured domains are shown as open-circles. Suppressor mutations identified in this study are colored magenta, and the residues mutated in the Dam1 phosphomimics are colored green. The marked residues are the closest approximations of the residues mutated in *S. cerevisiae*. N43 of Dad1 is conserved in *C. thermophilum*. The Dad2 residue K11 in *S. cerevisiae* is homologous to K35 in *C. thermophilum*. D119 of Spc34 maps to a short helix that contains H117 *C. thermophilum*. The bottom right shows the location of the Dad1 residue N43 on the surface of interface 2. Images were made using ChimeraX (Pettersen et al., 2021). (**B**) Left: representative images showing the localization of Dad3-mNeonGreen (Dam1c member) in preanaphase cells. Right: quantification of the intensity of Dad3-mNeonGreen at kinetochores/spindles in pre-anaphase cells. The amount of Dam1c localized to the spindle significantly reduced in cells with strong *bir1*Δ suppressors in the Dam1c. (**C**) Left: representative images showing the localization of Dad3-mNeonGreen in anaphase cells. Right: quantification of the intensity of Dad3-mNeonGreen at the spindle in anaphase cells. The intensity was measured in the middle of the spindle. (ns) non-signifcant; (*) P<0.05; (**) P<0.01; (***) P<0.001; (****) P<0.0001; unpaired t-test.

The oligomerization of the Dam1 complex into rings has been primarily implicated in microtubule binding (Miranda et al., 2005; Westermann et al., 2005). We therefore determined if the Dam1c mutants affect the localization of the complex to microtubules and kinetochores. We measured the amount of the Dam1 complex member Dad3 labeled with the fluorophore mNeonGreen at prometaphase and anaphase spindles in the presence of the suppressor mutants. The intensity of Dad3-mNeonGreen was significantly reduced at both prometaphase kinetochores/spindles and anaphase spindles for the three Dam1c suppressor mutations with the strongest rescue phenotypes (Figures 5B and 5C). The *dam1(S20D)* mutation also has a significant decrease in Dad3 localization in prometaphase. No significant decreases in Dad3 spindle localization were observed for *cdc4(G439S), mps1(V631M)*. Interestingly, the decrease in localization for the Dam1c mutants was much more pronounced along the anaphase spindle than at anaphase kinetochores, suggesting that the localization phenotypes are more likely to result from defects in microtubule binding than kinetochore association (Figure S3). We conclude that the Dam1c suppressor mutations decrease the amount of Dam1 complex at kinetochores and microtubules and this potentially occurs by decreasing the ability of the complex to oligomerize.

### Mps1 suppressor mutations do not act through previously established pathways

We next wanted to determine how the Mps1 mutations rescue Bir1 deletion. Mps1 kinase functions in activating the spindle assembly checkpoint at unattached kinetochores and recruiting the CPC to the inner centromere for chromosome biorientation (Weiss and Winey, 1996; van der Waal et al., 2012). In yeast, it also has an essential function in spindle pole body duplication (Winey et al., 1991). Disruption of any of these functions would be expected to decrease, rather than increase, the ability of cells to function with reduced CPC activity. One possible explanation for this would be if the Mps1 suppressor mutations have a gain-of-function phenotype that increases the kinase’s activity. We therefore tested if the Mps1, Cdc4, and Dam1c suppressors have gain-of-function activity by determining if they have a dominant phenotype in the heterozygous state. Only the Dam1c mutation *duo1(P17L)* had any improved growth when heterozygous, indicating that neither the Cdc4 nor the Mps1 mutations are gain-of-function (Figure S4A). To determine if the suppressor mutations affect the ability of Mps1 to activate the spindle assembly checkpoint, we measured the percentage of cells with Bub3-mNeonGreen foci either with or without the addition of nocodazole to depolymerize microtubules and create unattached kinetochores. The levels of Bub3 foci were unaffected by the *mps1(V631M)* mutation in either the presence or absence of nocodazole, demonstrating that the mutant cells are capable of activating the spindle assembly checkpoint (Figure S4B).

Since the Mps1 suppressor alleles have a recessive, partial loss-of-function phenotype and the mutations are located in the kinase domain, we next tested if a partial loss of Mps1 kinase activity can rescue Bir1 deletion. We used a mutation in the kinase domain that renders it sensitive to ATP analogs (Jones et al., 2005). This mutation, *mps1-as1*, improved the doubling time of *bir1Δ* cells to a similar extent to the suppressor mutation *mps1(V161M)* even in the absence of the small molecule inhibitor (Figure S4C). This result suggests that the analog-sensitive mutation partially decreases kinase activity on its own, as has been previously observed for similar alleles in other kinases (Bishop et al., 2000; Pinsky et al., 2006) and, further, that the Mps1 suppressor mutations affect its kinase activity to a comparable extent. Addition of higher concentrations analog inhibitor decreased growth in *mps1-as1*, *bir1Δ* cells, demonstrating that too little Mps1 activity is detrimental to growth even in the absence of Bir1.

We next tested if the Mps1 mutations are suppressing CPC activity though its phosphorylation of the Dam1 complex subunit Dam1. Mutations that prevent phosphorylation of Dam1 at S218 and S221 by Mps1 kinase have previously been demonstrated to decrease kinetochore-microtubule stability and partially rescue temperature-sensitive Ipl1 mutations (Shimogawa et al., 2006; 2010). However, we did not observe any rescue of Bir1 deletion with these mutants (Figure S4D). This result agrees with our experiments demonstrating that the Mps1 and Dam1c suppressor mutations act through different mechanisms (Figures 3A-C). We conclude that partial disruption of Mps1 kinase activity results in rescue of Bir1 deletion through a currently unknown mechanism.

### The SCF^Cdc4^ complex affects CPC localization to the spindle/kinetochores in prometaphase in yeast and human cells

Finally, we wanted to determine how the SCF mutations rescue Bir1 deletion. We first tested if a previously identified temperature sensitive mutation in *CDC4, cdc4-1*, is capable of rescuing loss of Bir1. We also tested a temperature-sensitive version of another F-box protein, Met30. Both Cdc4 and Met30 have recently been shown to affect the stability of the yeast CENP-A homolog Cse4 (Au et al., 2020). The *cdc4-1* mutation shows a degree of rescue nearly as strong as the suppressor mutation *cdc4(G439S)* (Figure 6A). In contrast, a temperature-sensitive mutation in Met30 failed to show any rescue of Bir1 deletion, demonstrating the specificity of the suppression phenotype for SCF^Cdc4^. We conclude that rescue of *bir1Δ* results from decreased SCF^Cdc4^ activity and is not unique to the mutations identified in our screen.

**Figure 6.**
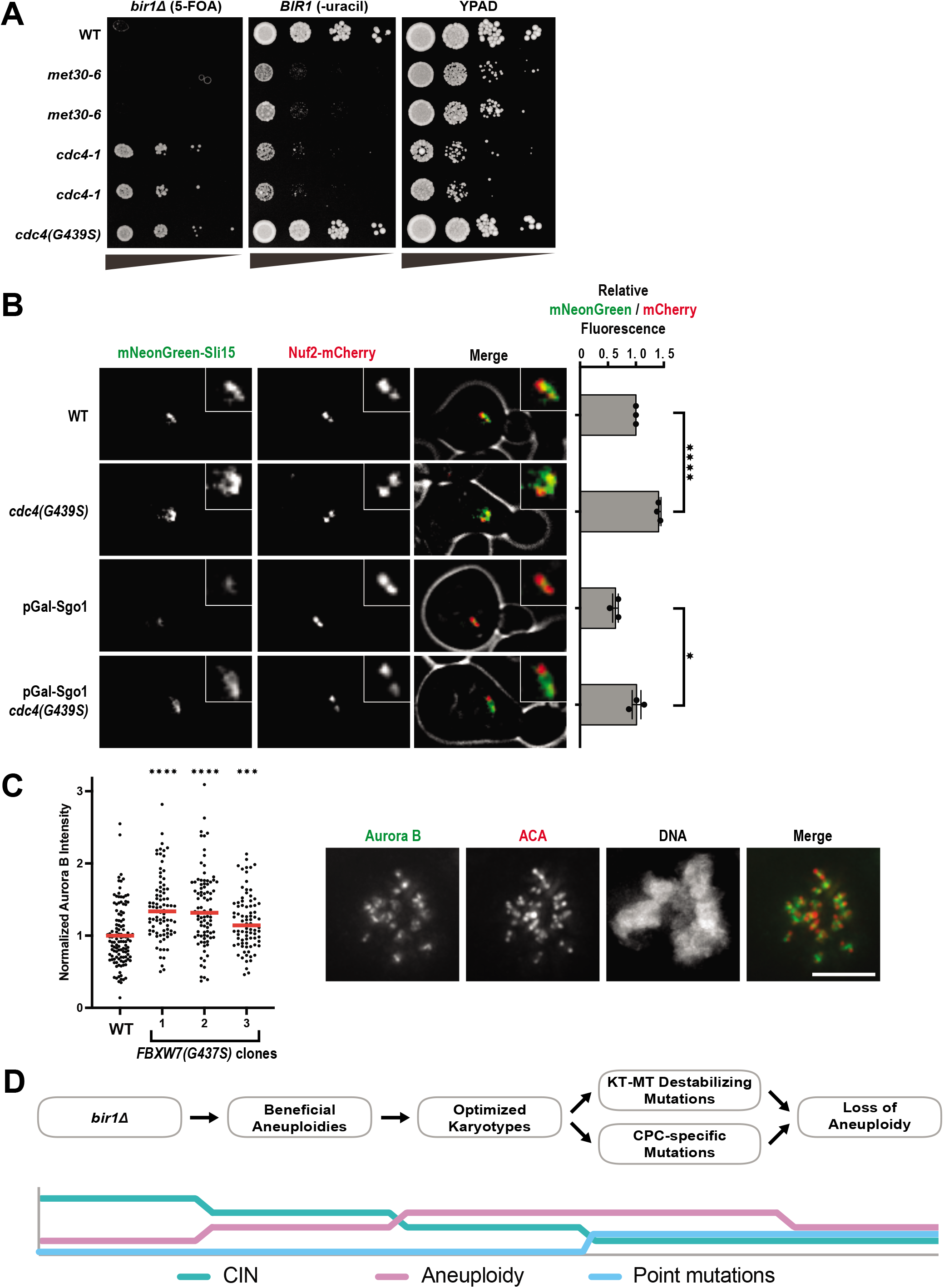
Cdc4 limits CPC localization in prometaphase independently of Sgo1. (**A**) Serial dilutions of strains engineered with the indicated mutations were tested for rescue of Bir1 deletion. 10-fold serial dilutions on the indicated media are shown. The *met30-6* and *cdc4-1* alleles are temperature sensitive. Two independent clones of each of these alleles are shown. The serial dilutions were performed at the permissive temperature (20°C). (**B**) The amount of mNeonGreen-Sli15 (CPC member) located at kinetochores (Nuf2-mCherry) was measured in prometaphase. Sgo1 was depleted by placing it under a galactose-inducible promoter and switching to glucose-containing media (YPAD). Representative images are on the left. (**C**) Measurements of Aurora B intensity at the inner centromere of prometaphase kinetochores. Image from an example WT cell is shown to the right. Scale bar is 5 *μ* m long. Each data point is an individual chromosome. More than 15 cells for each cell line were measured across two independent experiments. (**D**) Timeline of adaptation to high levels of CIN through aneuploidy and point mutations. (ns) non-signifcant; (*) P<0.05; (***) P<0.001; (****) P<0.0001; unpaired t-test.

To determine how Cdc4 activity affects CPC function, we first determined if the ubiquitin ligase directly degrades the CPC subunits Sli15 or Ipl1. We monitored Sli15 and Ipl1 protein levels in cells that were prevented from expressing new protein by the addition of cycloheximide. Sli15 and Ipl1 protein levels were unchanged by the presence of the *cdc4(G439S)* suppressor mutation, indicating that this mutation does not rescue Bir1 deletion by increasing the levels of Sli15 or Ipl1 (Figure S5). We next tested if the Cdc4 suppressor mutations change the localization of the CPC. *cdc4(G439S)* significantly increased CPC localization to prometaphase spindles/kinetochores by ~50% (Figure 6B). We wanted to determine if this mutant also rescues CPC localization in *bir1Δ* cells. However, *bir1Δ* greatly reduces Sli15 expression (Campbell and Desai, 2013), so we instead determined if *cdc4(G439S)* rescues localization after depletion of Sgo1, the upstream recruiter of Bir1 to the inner centromere (Figure 1A). mNeonGreen-Sli15 localization was significantly increased in prometaphase after Sgo1 depletion, demonstrating that this increase in CPC localization is independent of the Sgo1 recruitment pathway that acts through Bir1 (Figure 6B).

We next tested if mutations in the human homolog of Cdc4, FBXW7, also affect the accumulation of the CPC at the inner centromere. We therefore engineered the sole copy of FBXW7 in the haploid HAP1 chronic myeloid leukemia cell line with the equivalent of the G439S point mutation in the WD-40 domain that we identified in adapted *bir1Δ* cells (Figures S6A and S6B). Of note, mutations in this region of the protein are frequently observed in uterine and colon cancer (Yeh et al., 2018). This mutation (G437S in humans) resulted in a slight increase in colony size, consistent with the function of FBXW7 as a suppressor of cell cycle entry (Figure S6C). Intriguingly, all three cell lines engineered with this mutation showed an increase in Aurora B staining at the inner centromere in prometaphase (Figure 6C). This ~50% increase is similar to what we observe in yeast, indicating that this function of the SCF complex in CPC localization is conserved. We conclude that SCF^Cdc4/FBXW7^ activity limits the recruitment of the CPC to the spindle and/or kinetochores, and that reduction of this function partially restores CPC localization when the inner centromere recruitment pathway is disrupted.

## Discussion

In cancers, cells with high levels of CIN and aneuploidy often have overstabilized kinetochore-microtubule connections (Bakhoum et al., 2009a). Although the mechanisms that cause this phenotype are still largely unclear, one way to decrease microtubule turnover at the kinetochore is through decreased CPC activity (Cimini et al., 2006). In this study, we identify multiple mechanisms that cells use to adapt to the overstabilization of microtubules resulting from the deletion of the CPC subunit Bir1/Survivin. We determined that *bir1Δ* suppressor mutations act through two distinct mechanisms. The first mechanism involves mutations in proteins that directly attach the kinetochore to microtubules; such mutations were identified in five out of ten subunits of the Dam1 complex. The second mechanism results from mutations in genes that appear to affect the CPC more directly, which includes members of the SCF complex, the kinase Mps1, and the CPC member Sli15.

All of the suppressor mutations that we identified decrease the levels of CIN that result from Bir1 deletion. Notably, we did not identify mutations whose function suggests that they allow for the greater tolerance of aneuploidy. Furthermore, a mutation in the gene *UBP6* that has been previously demonstrated to reduce the negative effects of aneuploidy did not rescue Bir1 deletion (Torres et al., 2010). However, mutation of *UBP6* was previously shown to improve the growth of only a subset of aneuploid chromosomes (5, 8, 9, and 11). Of these chromosomes, only aneuploidy of chromosome 8 is frequently observed in strains adapted to *bir1Δ*. Therefore, aneuploidy-tolerating mutations could potentially improve the growth of cells with other sources of CIN that adapt through different aneuploid chromosomes.

In our adaptation experiments, we identified mutations in three different kinetochore complexes that directly bind to microtubules. The majority of these mutations were in the Dam1 complex. These mutations decrease the localization of the complex to microtubules and likely interfere with the complex’s ability to oligomerize. The cellular phenotypes that we observe for these mutations are consistent with increased turnover of kinetochore microtubules. In vertebrates, kinetochore microtubule turnover can be increased through the overactivation of the kinetochore-localized kinesins MCAK and Kif2b (Walczak et al., 1996; Kline-Smith and Walczak, 2002; Bakhoum et al., 2009b). Overexpression of either of these kinases has been shown to decrease the mitotic errors observed in some cancer cell lines (Bakhoum et al., 2009b). Furthermore, adaptation of human cells to a drug that activates MCAK resulted in cells with decreased Aurora B activity (Orr et al., 2016). Together, these results all point to the importance of a balance between Aurora B activity and other factors that affect the stability of connections between kinetochores and microtubules.

Mps1 is a highly conserved kinase that functions in sensing unattached kinetochores. Mps1 phosphorylates Spc105/Knl1 on MELT repeats, which recruits the spindle assembly checkpoint proteins Bub1 and Bub3/BubR1. CPC mutations have been previously shown to have synthetic growth defects with spindle assembly checkpoint mutations, indicating that decreased SAC activity does not generally rescue CPC mutants (Ng et al., 2009). Bub1 binding to Spc105/Knl1 also contributes to the localization of the CPC to the inner centromere, which is one of the key locations for its function in chromosome biorientation. Bir1 is required for the CPC to bind to the inner centromere, so this targeting function of Mps1 should not affect cells with Bir1 deletion (Shimogawa et al., 2009; Makrantoni and Stark, 2009). In budding yeast, Mps1 has an additional essential function in spindle pole body duplication. Interestingly, we would predict that decreased activity of any of these three functions would make the *bir1Δ* phenotype worse, not better. We conclude that the suppressor mutations partially decrease kinase activity and rescue via an unidentified mechanism. In a large-scale screen for synthetic interactions between temperature-sensitive mutations, it was found that some combinations of *MPS1* and *IPL1* alleles had positive genetic interactions, while others had negative interactions (Costanzo et al., 2016). These results support the idea that a balance between Mps1 activity and CPC activity is required for accurate chromosome segregation. Additional research will be required to determine which specific functions of Mps1 and which of its substrates are responsible for rescuing Bir1 deletion.

The SCF ubiquitin ligase complex is a key regulator of many pathways related to cell cycle entry. It degrades factors that inhibit cyclins, allowing for the activation of cyclin-dependent kinases. The human homolog of the SCF F-box protein Cdc4 is FBXW7/hCDC4. FBXW7 is a strong tumor suppressor that is frequently mutated in many cancer types (Spruck et al., 2002). This function is largely attributed to the ability of FBXW7 to target important oncogenes for degradation, including MYC and Cyclin E (reviewed in Yeh et al., 2018). In addition to its roles in regulating cell cycle initiation, FBXW7/Cdc4 also has functions in mitosis. In yeast, certain Cdc4 mutants arrest in mitosis prior to anaphase (Goh and Surana, 1999). More recently, it was shown that Cdc4 contributes to the degradation of the inner kinetochore protein Ame1 (Böhm et al., 2021). In human cells, FBXW7 mutants are sensitive to inhibitors of the spindle assembly checkpoint, demonstrating a potential function in mitosis (Bailey et al., 2015). However, the mechanisms by which SCF^Cdc4^ regulates mitosis are still unclear. Our screen for *bir1Δ* suppressors has uncovered a connection between the SCF^Cdc4^ complex and regulation of the CPC in budding yeast. These suppressor mutations in the SCF complex increase the localization of the CPC to the spindle/chromosomes in the absence of inner centromere targeting, which suggests that the SCF^Cdc4^ complex may play a role in regulating alternative mechanisms of CPC localization in early mitosis.

Determining how cells adapt to CIN caused by the overstabilization of kinetochore-bound microtubules has important implications in cancer. Additionally, Aurora kinase inhibitors are actively being investigated in a variety of combination therapies in both pre-clinical and clinical trials (reviewed in Du et al., 2021). It is therefore important to know how cells adapt to the overstabilization of kinetochore attachments. Here, we have outlined a timeline of events that cells use to adapt to decreased Aurora B activity (Figure 6D). First, cells obtain specific aneuploidies that partially decrease CIN. Next, the aneuploid karyotypes are refined to obtain an optimal complement of aneuploid chromosomes (Ravichandran et al., 2018). Following adaptation through aneuploidy, specific point mutations are acquired that further decrease the rate of CIN. These point mutations can affect the CPC itself or independently increase the turnover of kinetochore microtubules. Finally, point mutations that reduce CIN in a more targeted way allow for a decrease in the number of aneuploid chromosomes.

This timeline of adaptation supports the theory that aneuploidy often provides a rapid but temporary form of adaptation, as has previously been observed in yeast adapted to heat stress or a tubulin mutation (Yona et al., 2012; Pavani et al., 2021). In both of those studies, a single chromosome was gained early in the adaptation process and sometimes lost again at later time points. Here, we observe the reversion towards the euploid state for many different chromosomes resulting from a wide variety of mutations. This return to the original copy number could help explain the high prevalence of whole chromosome loss of heterozygosity (LOH) in cancers. In colorectal cancers for example, whole chromosome LOH is extremely common, even though the chromosomes are often present in multiple copies (Thiagalingam et al., 2001). This theory could help explain why the bulk of LOH events do not contain known tumor suppressor genes, as the LOH itself was not the driver event, but simply a byproduct of temporary adaptation through aneuploidy (Ryland et al., 2015).

## Materials and methods

### Yeast strains and media

All yeast strains and plasmids that were used in this study are listed in Supplemental Table S1. Strains were grown in yeast extract and peptone supplemented with 40 μg/mL adenine-HCl (YPA) and sugars (2% glucose: YPAD, 1% galactose and 1% raffinose: YPAGR). Benomyl (Sigma-Aldrich, 381586), nocodazole (VWR, 487928) and 5-FOA (Chempur, 220141-70-8) were used at concentrations of 10 μg/mL, 15μg/mL. and 1 mg/mL, respectively. All cultures were incubated at 30°C unless otherwise stated. Gene deletions were carried out as previously described (Longtine et al., 1998). The engineering of specific mutations at endogenous loci was achieved using the method established by the Boone lab (Li et al., 2011).

### Tissue Culture

All cell lines in this study tested negatively for mycoplasma contamination. HAP1 cell lines were cultured in a humidified growth chamber at 37°C and 5% CO_2_ in Iscove’s Modified Dulbecco’s Medium (IMDM) (Sigma-Aldrich) supplemented with 10% Fetal Bovine Serum (Thermo Fisher Scientific) and 1% (v/v) Penicillin-Streptomycin (Sigma-Aldrich).

For mutation of the endogenous FBXW7 locus in HAP1 p53^-^ cells, a CRISPR/Cas9 strategy was applied. SgRNAs were cloned into pSpCas9(BB)-2A-GFP (PX458, Addgene plasmid #48138). For homologous recombination, a repair template carrying the respective mutation and 1000 bp homology flanks was synthesized as gBlock gene fragment (Integrated DNA Technologies IDT) and inserted into the plasmid pmScarlet_C1 (Addgene plasmid #85042). The plasmid mix of guide RNA plasmid and the repair template was transfected into HAP1 cells using FuGENE HD (Promega). Two days after transfection, cells were sorted for the presence of Cas9 (GFP positive) and repair template (mScarlet positive). Three days later, single cells were sorted into 96 well plates. Clonal populations were expanded gradually over the course of three weeks. The FBXW7 mutations were identified by Sanger Sequencing and karyotypes of the cell lines were validated by flow cytometry and whole-genome sequencing.

### Flow Cytometry

HAP1 cells were trypsinized and stained with 10μg x ml^-1^ Hoechst 33342 (Thermo Fisher Scientific) for 30min at 37°C. Cells were then analyzed for their ploidy state (based on G1 haploid peaks) in PBS supplemented with 5% FBS in a FACSAria III (BD) flow cytometer.

### Next-generation sequencing and data analysis

All protocols for genomic DNA isolation, library preparation, and sequencing data analysis used here were previously described by Ravichandran et al (Ravichandran et al., 2018). GOrilla gene-ontology tool (http://cbl-gorilla.cs.technion.ac.il/) was used for GO enrichment tests. The target gene list was composed of all genes with non-synonymous mutations in the adapted strains (gene repeats included), and the background list was all 6002 genes annotated in the *S. cerevisiae* genome.

### Microscopy

Strains were grown overnight in YPAD or YPAGR, subsequently diluted 1000-fold in the morning into fresh media, and grown for 4-6 h to mid-log phase. Cells were pelleted by brief centrifugation, washed twice with 1 mL water, and resuspended in 200 μL water. 2 μL of cell suspension was spotted onto 1% agarose pads supplemented with complete synthetic media (SC) and 2% glucose. A coverslip was placed on top and sealed with 1:1:1 mixture by weight of paraffin (Merck), lanolin (Alfa Aesar) and Vaseline (Ferd. Eimermacher). Images were collected on a DeltaVision Ultra Epifluorescence Microscope system (Cytiva) at 30°C and a PlanApo N 60/1.42 Oil objective and a sCMOS sensor, 1020 x 1020 pixels, 6.5 μm pixel size camera. For live yeast imaging, 12 z-sections with step size of 0.5 μm were taken. Images were deconvolved using softWoRx software (Life Sciences Software). Counting of Bub3 foci was done using deconvolved images. Quantification of Dad3 fluorescence intensity was performed on non-deconvolved images using ImageJ and a custom-made Python script (Fink et al., 2017). All analyses were obtained from images obtained from a minimum of three different days. Representative images are deconvolved and were contrast adjusted identically using ImageJ.

To test the localization of Sli15 to the mitotic spindle, endogenous Sli15 was put under the control of a *Gal10-1* promoter. For synchronization and depletion of endogenous Sli15, overnight cultures grown in YPAGR were diluted 1:100 into YPAGR and shaken for 2h at 30°C. Cells were resuspended in YPAGR containing α-factor (10 μg/mL) for 45 min at 30°C. Subsequently, the medium was exchanged with YPAD containing α-factor (10 μg/mL) and grown for 2h and 15 min at 30°C. Cells were then released from the G1 arrest into the cell cycle by washing twice with YPAD and resuspending in YPAD containing Pronase E (Merck). Synchronized yeast cells were released into the cell cycle and grown for 45 min at 30°C. The cells were then pelleted, washed with 1 mL sterile water, resuspended in 100 μL sterile water, and 2 μL cell suspension was put onto 1% agarose SC pads.

For the immunofluorescence of human cells, mitotic cells were collected by mechanical shake-off and immobilized on adhesion slides (Marienfeld) for 30min at 37°C. Next, cells were washed once in PBS, fixed for 15min with 4% Formaldehyde in PBS and subsequently permeabilized using 0.5% Triton-X-100 in PBS (0.5% PBST). Cells were blocked for 1h in 0.01% PBST + 2% Bovine Serum Albumin and co-stained overnight with rabbit monoclonal anti-Aurora B antibody (Abcam, ab45145, 1:200) and human anti-centromere antibody (Antibodies Incorporated, 15-234, 1:200). After several washes with 0.01% PBST, cells were co-stained with goat anti-rabbit IgG Alexa Fluor 488 (Thermo Fisher Scientific, A32723, 1:500) and goat anti-human IgG Alexa Fluor 647 (Invitrogen, A-21445, 1:500) for 2 hours. After Immunostaining, DNA was stained for 1min with 1μg/ml DAPI (Thermo Fisher Scientific).

### Minichromosome loss assay

Yeast strains transformed with the minichromosome pCC250 were grown to saturation overnight in SC-uracil media, diluted to a starting OD^600^ of 0.05, and grown for 24 hours in 2mL of YPAD media. After 24 hours of growth, these strains were diluted 10^6^-fold in sterile water and 100μL of each strain was plated onto an SC-uracil plate and a YPAD plate. The plates were incubated at 30°C for 48 hours. Pictures taken of the subsequent colonies by a Canon EOS Digital Camera. The colony numbers were counted using the “Analyse Particles” function in ImageJ. The number of colonies on the *ura-* plates were then divided by the number that grew on the YPAD plate to obtain the measurement of transmission fidelity.

### ATP-analog growth assay

Strains were grown overnight in YPAD and diluted to a starting OD^600^ of 0.05 in 3mL YPAD at 25°C. The strains were given up to 4.5 hours to reach exponential phase. At time point zero, the indicated amount of ATP-analog 1NM-PP1 (Merck) or DMSO was added to the strains. Immediately after drug addition, 0.5mL of each sample was diluted 2x in a 1mL cuvette to analyse the OD^600^ (timepoint 0). This was repeated every 1.5 hours thereafter, for 7.5 hours.

### Budding index

Strains were grown overnight in YPAD media, and 100μL of each overnight was added to 2mL YPAD. After allowing 1 hour for recovery, 2μL of α-factor was added every 45 minutes for 2 hours. Strains were then washed using fresh YPAD media mixed with Pronase E (Merck) (1:10). The number of large-budded cells was counted via light-microscopy every half an hour for 2.5 hours. 100 cells were counted per slide per time-point.

### Colony Formation Assay

On day one, 600 cells were seeded into 6-well plates. Drugs were added at the indicated concentrations and the plates were incubated for 12 days. Colonies were then fixed with 4% Paraformaldehyde in PBS (Sigma-Aldrich) for 20min, washed with water, stained for 30min with Crystal Violet, washed with water, and dried.

### Protein extraction and Western blotting

For yeast protein extraction, saturated overnight cultures were diluted in 30 mL YPAD in order to obtain an OD^600^ of 0.25. Cells were grown at 30°C while shaking until an OD^600^ of 0.7 was reached. At this point cycloheximide (50 μg/mL) was added (t = 0). 5 mL aliquots were taken every 30 min for a 2h period at 30°C. Immediately after each aliquot was taken, proteins were extracted by pelleting the cells and resuspending them in 100 μL 5% trichloroacetic acid. Following 10 min incubation at room temperature, the cells were washed once with 1 mL ddH2O and resuspended in 100 μL lysis buffer (50 mM Tris, pH 7.4, 50 mM dithiotreitol, 1 mM EDTA, Complete EDTA-free protease inhibitor cocktail (Roche) and Phosstop (Roche)). Cells were vortexed for 30 min at 4°C after the addition of glass beads. Subsequently, 33 μL 4x sample buffer was added and incubated at 95°C for 5 min. Samples were stored at −20°C. For immunoblots, the following antibodies were used: rat anti-HA-clone 3F10 (Roche), mouse anti-PGK1 monoclonal 22C5D8 (Thermo Fisher Scientific). Membranes were probed with the corresponding secondary antibodies: anti-rat IgG-HRP-lined (Cell Signaling Technology) or anti-mouse IgG-HRP-linked (Cell Signalling Technology). Immunoblots were quantified using ImageJ.

### Statistics

Statistical analysis was performed using GraphPad Prism software. Details to the statistical tests used in a particular experiment are reported in the figure legends.

## Acknowledgements

The authors thank the Campbell and Dammermann laboratories for helpful discussions and comments. We thank the Basrai and Klein labs for gifts of yeast strains and plasmids. The parental HAP1 p53-null cell line was kindly provided by J.I. Loizou. We acknowledge the service of the Perutz Labs BioOptics Facility, a member of the VLSI. We would also like to acknowledge the services provided to us by the VBC sequencing facility located at the VBCF.

This work was supported by Vienna Science and Technology Fund (WWTF) grant VRG14-001 and Austrian Science Fund (FWF) grants Y944-B28 and W1238-B20 to C.S.C.

## Author contributions

T.M. performed the experiments in Figure 6B and S5. M.C.R. performed experiments for Figure 1C and D. M.A.Y.A. performed the experiments for Figures 6C and S6. M.N.C. performed the rest of the experiments. T.M., M.N.C. and C.S.C. conducted data analysis. C.S.C. and M.N.C. wrote the manuscript. C.S.C. supervised the project.

## Supplementary figure legends

**Figure S1. CIN-related mutations in *bir1*Δ-*ad2* strains are associated with decreased aneuploidy.**

(**A**) Histogram showing the proportion of strains that are aneuploid for specific chromosomes in partially adapted (*bir1*Δ-*ad*) and further adapted (*bir1*Δ-*ad2*) strains. There is less aneuploidy overall in the *bir1*Δ-*ad2* strains, but not every chromosome is decreased in frequency. (**B**) 874 genes out of 6002 total yeast genes (14.5%) have phenotypes related to CIN. The relative number of identified mutations in the *bir1*Δ-*ad2* and *bir1*Δ-*ad* strain collections is shown. (**C**) Serial dilutions of strains engineered with the *ubp6(E256X)* aneuploidy tolerance mutation with and without selection for Bir1 deletion. 10-fold serial dilutions on the indicated media are shown. *ubp6(E256X)* does not suppress *bir1*Δ. (**D**) Histogram showing the proportion of strains that are aneuploid for specific chromosomes in *bir1*Δ-*ad2* strains that have identified suppressor mutations, and those that do not.

**Figure S2. Double mutants of *bir1*Δ suppressors in the Dam1 complex and the *dam1(S20D)* phosphomimic have strong synthetic phenotypes**

(**A**) Cell cycle progression as measured by percentage of large budded cells over time after release from G1 arrest. Wild-type (WT) cells treated with nocodazole and benomyl were used as a positive control for mitotic arrest. None of the tested suppressor mutants cause a mitotic delay. (**B**) Tetrad dissections of diploids heterozygous for both a *bir1*Δ suppressor mutant and the *dam1(S20D)* phosphomimic mutant. Presence of either the suppressor mutant or *dam1(S20D)* was determined by antibiotic resistance conferred by a gene intergrated downstream of the mutations (clonNAT and G418 respectively). Spore viability was calculated as the proportion of colonies that grew relative to the total colonies that could have grown. (**C**) Cell cycle progression as measured by percentage of large budded cells over time after released from G1 arrest. Rare surviving *duo1(P17L), dam1(S20D)* double mutants have a strong mitotic delay. (**D**) Proportion of colonies that grow on plates with restrictive (lacking uracil) versus permissive (YPAD) media after 48 hours growth under permissive conditions with a (*URA3*-containing plasmid. *duo1(P17L), dam1(S20D)* double mutants have a strong decrease in transmission fidelity. (ns) not significant; (*) P<0.05; (**) P<0.01; unpaired t-test.

**Figure S3. Dam1 complex localization to spindles and kinetochores in anaphase *bir1*Δ suppressors in the Dam1 complex.**

Quantification of the intensity of Dad3-mNeonGreen at kinetochores in anaphase cells. The differences in localization at the anaphase kinetochores are much weaker than those measured along the spindle (Figure 5C).

**Figure S4. Suppression of *bir1*Δ by Mps1 mutations occurs through decreased kinase activity that does not inhibit spindle assembly checkpoint activity.**

(**A**) Serial dilutions of strains engineered with the indicated mutations were tested for rescue of Bir1 deletion. 10-fold serial dilutions on the indicated media are shown. Diploid strains were heterozygous for either an identified suppressor mutation or a full deletion of the gene. (**B**) Quantification of the proportion of prometaphase cells with Bub3-mNeonGreen foci with or without 15μg/mL nocodazole. The *mps1(V631M)* does not affect the percentage of cells with spindle assembly checkpoint foci. (**C**) Growth assays conducted with an ATP-analog (1NM-PP1) that specifically inhibits the *mps1-as1* allele. Strains with wild-type Mps1 were not affected by the inhibitor (left). (**D**) Serial dilutions of strains engineered with the indicated mutations were tested for rescue of Bir1 deletion. 10-fold serial dilutions on the indicated media are shown. Dam1 alleles with nonphosphorylatable mutations in Mps1 phosphosites (S218 and S221) do not rescue growth after *bir1*Δ.

**Figure S5. *bir1*Δ suppressor mutations in Cdc4 do not affect protein stability of Sli15 or Ipl1.**

Western-blot of a time series of Ipl1 and Sli15 protein levels after inhibition of protein translation with cycloheximide. Cycloheximide was added to cells at timepoint 0, and cells were harvested and fixed at 30 minute intervals. Representative images from one experiment are shown on top and quantification of three independent experiments is shown on bottom. Error bars represent standard-deviation of three biological replicates from Western Blot quantifications. There is no detectable change in the degradation of Sli15 or Ipl1 with or without the *cdc4(G439S)* mutant.

**Figure S6. Engineering and characterization of FBXW7 mutant cell lines.**

(**A**) Strategy implemented to introduce specific point mutations in FBXW7 in HAP1 cells. Transfected cell lines were originally screened for a silent mutation that creates an EcoR1 restriction site. (**B**) Sanger sequencing confirms the presence of the G437S mutation in FBXW7. (**C**) Cell lines mutant for FBXW7 have an increased colony size after 12 days of growth in comparison to a cell line with wild-type FBXW7. Mean and standard deviations are shown. (***) P<0.001; (****) P<0.0001; unpaired t-test.

