## Supplementary figures and images for "Multiple routes of adaptation to high levels of CIN and aneuploidy in budding yeast"

### Supplemental Figures

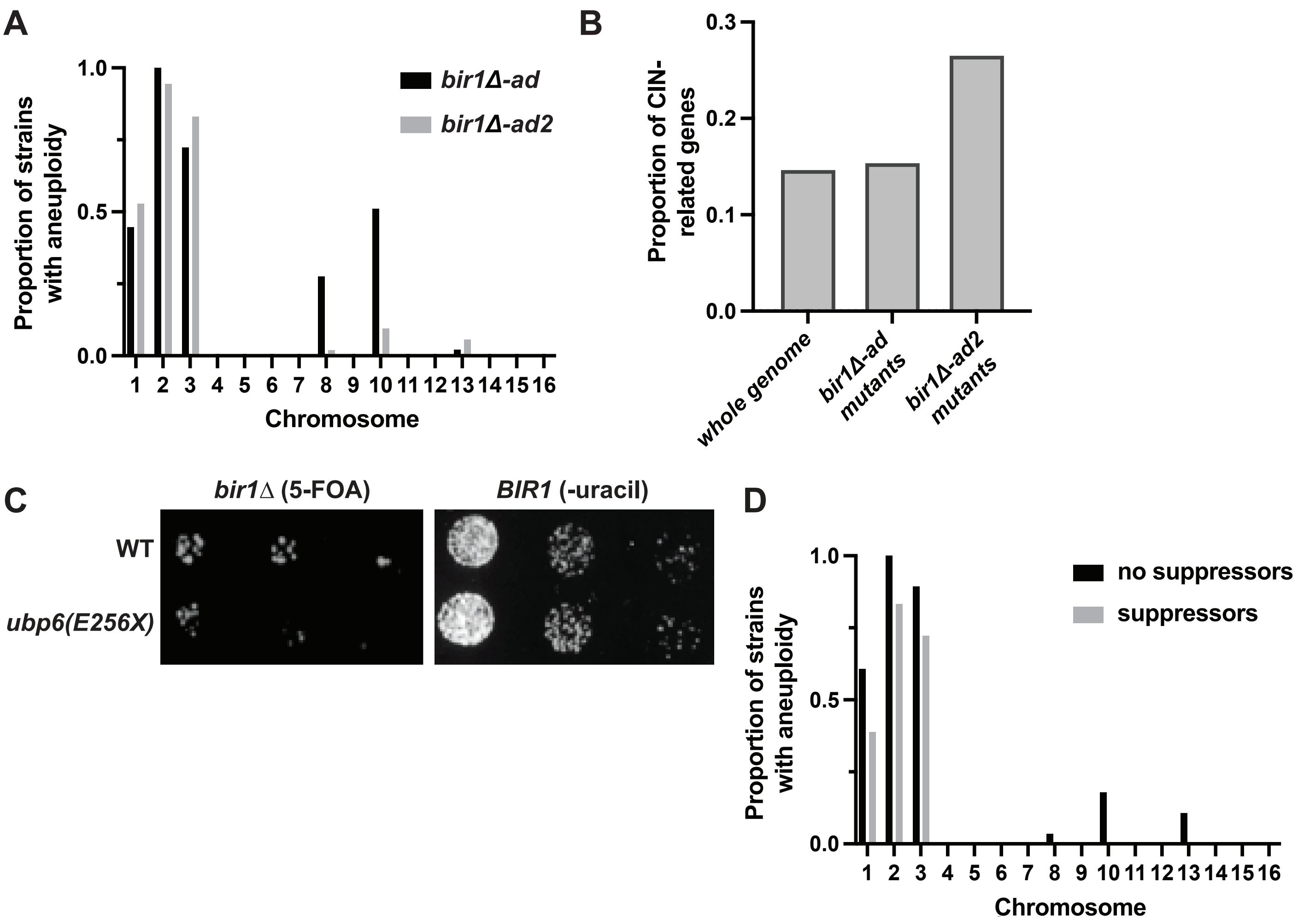

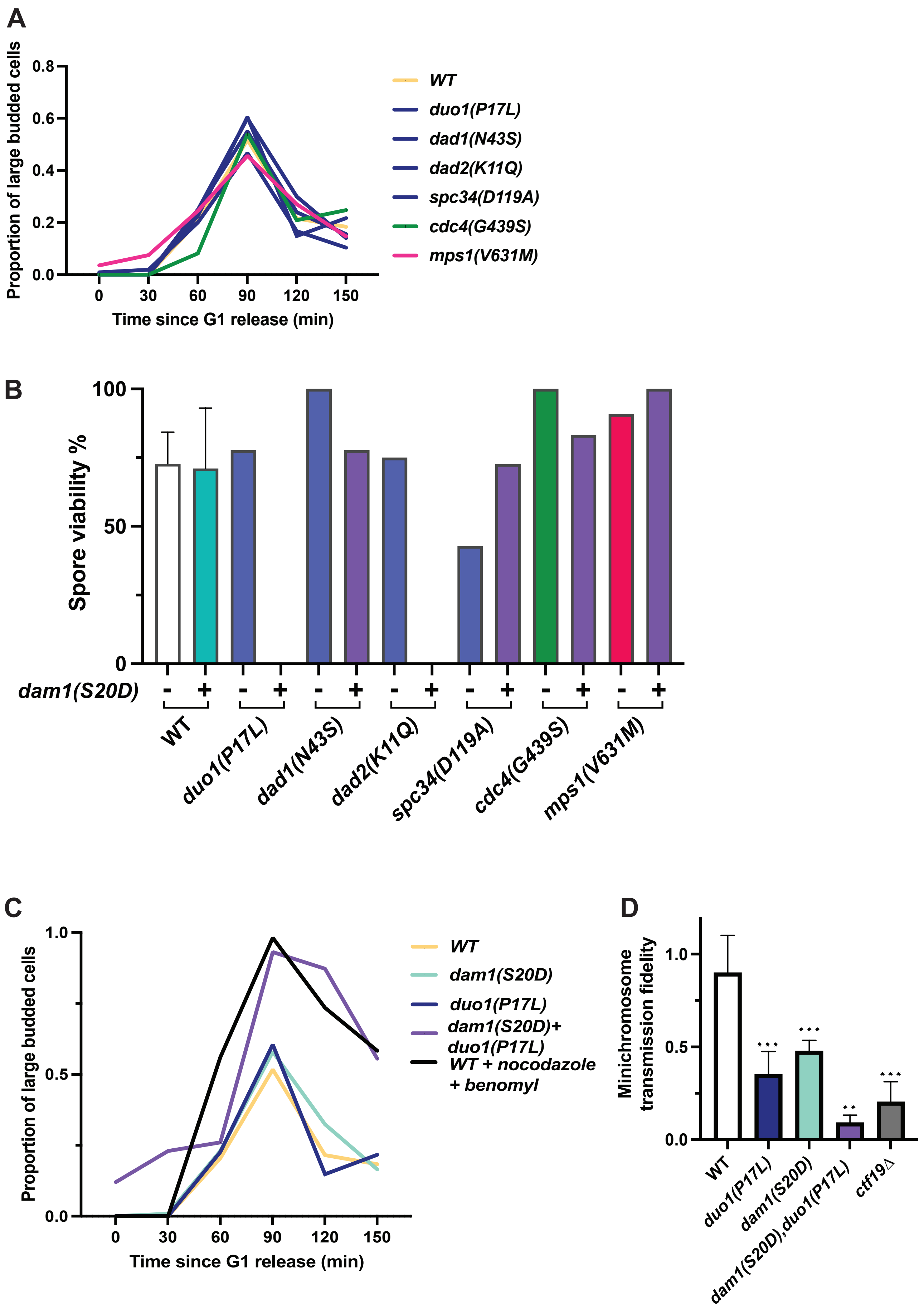

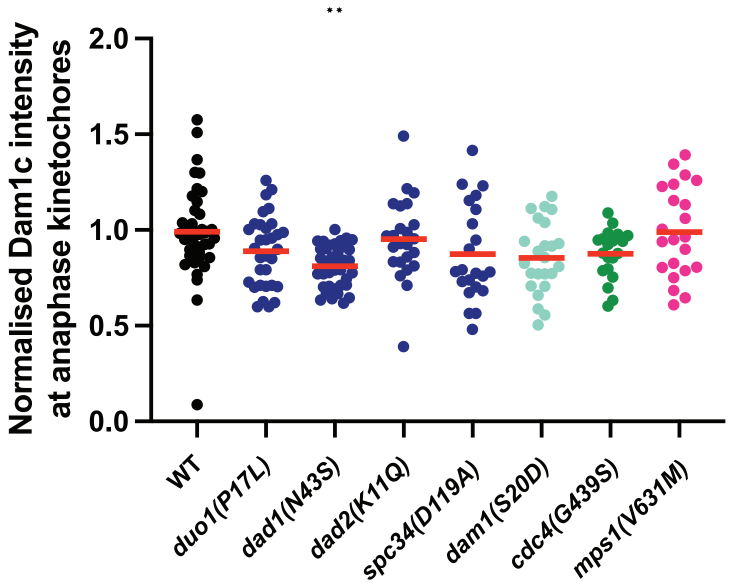

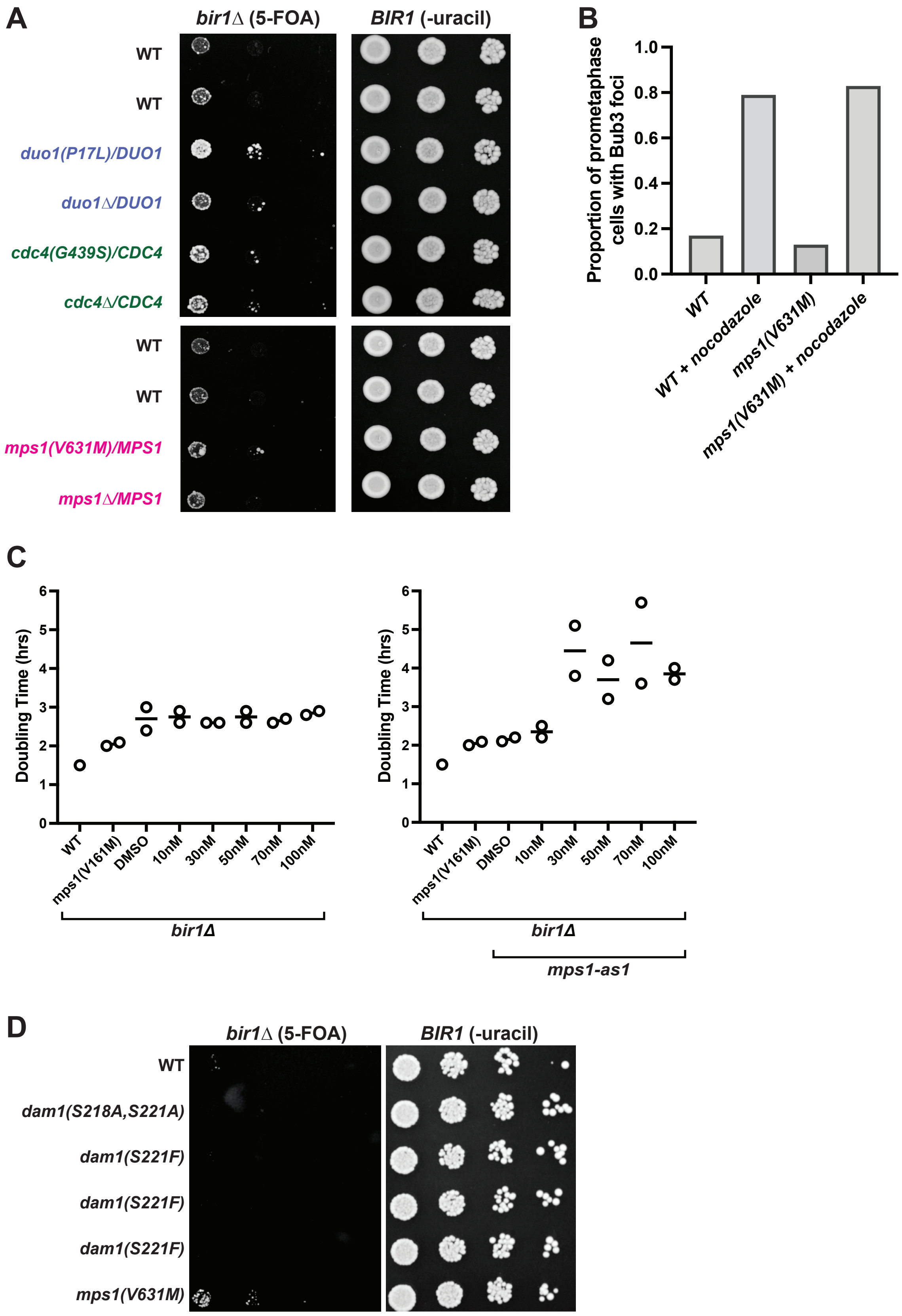

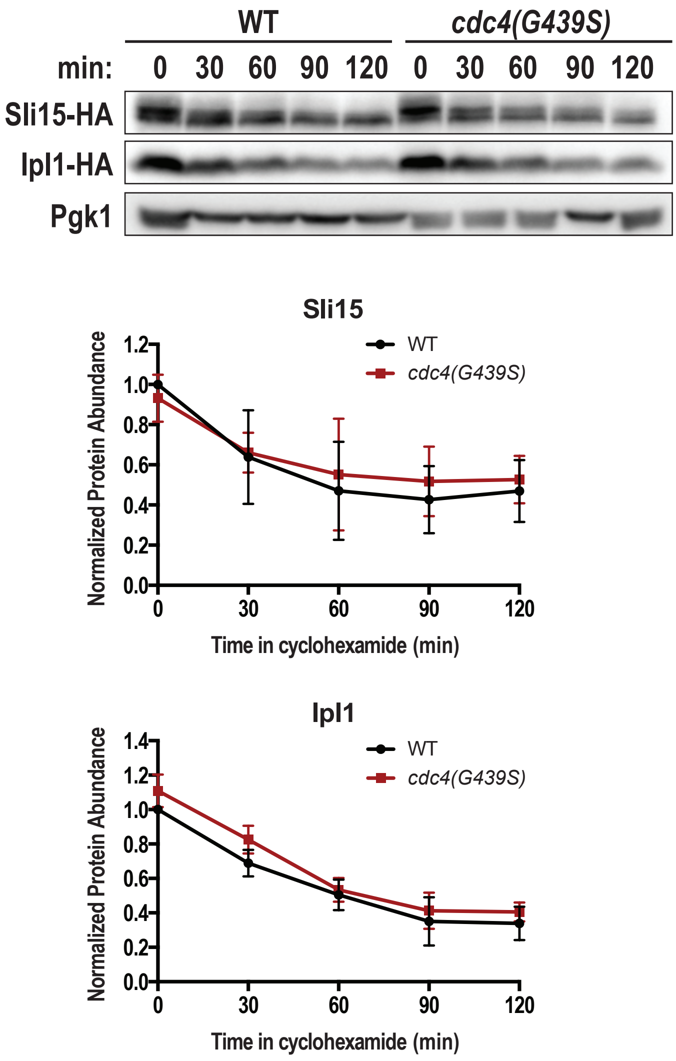

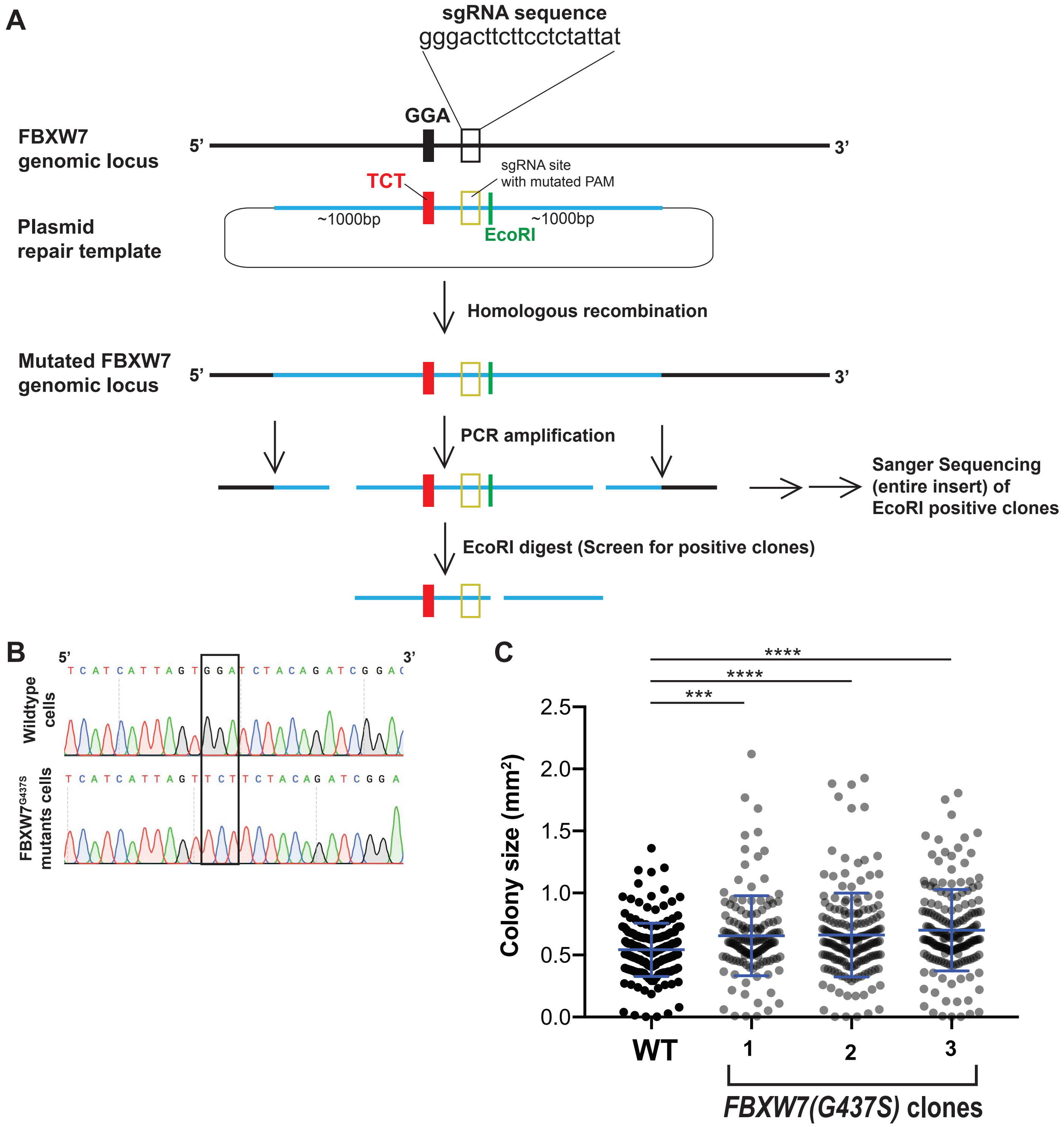
