## Supplemental Table 1 for "Multiple routes of adaptation to high levels of CIN and aneuploidy in budding yeast"

### Table S1. Yeast strains, Plasmids, and human cell lines from this study

| Strain | Genotype | Source | Background | Figure/s |
|---|---|---|---|---|
| CCY149 | MATa/α; Bir1+/Δ::HYG; ura3-1; LEU2/leu2,3-112; his3-11:pCUP1-GFP12-lacI12:HIS3 trp1-1:256lacO:TRP1; LYS2/lys2Δ ADE2/ade2-1; can1-100; bar1Δ | a | W303 | 1; 2; S1 |
| *bir1Δ* haploids | These strains were made either through tetrad dissection of CCY149 or from single colonies of CCY1905 selected on FOA plates. | c | W303 | 1B;1C;1D |
| *bir1Δ-ad* haploids | All strains were derived from tetrad dissecting CCY149 (102 haploid strains) | b | W303 | 1B;1C;1D;1E;2F;S1A;S1B;S1D |
| *bir1Δ-ad2* haploids | All strains were derived from bir1Δ-ad (68 strains) | b | W303 | 1B;1C;1D;1E;2D;2E;2F;S1A;S1B;S1D |
| *bir1Δ-ad3* haploids | All strains were derived from bir1Δ-ad2 (15 strains) | c | W303 | 1B;1C;1D;1E;2D;2E;2F;S1A;S1B;S1D |
| CCY434 | MATa; ura3-1; leu2;3-112; his3-11; trp1-1; ade2-1 | a | W303 | 3A; 4D; S2A; S2C |
| CCY747 | MATα; his3-11:pCUP1-GFP12-lacI12:HIS3 trp1-1:256lacO:TRP1; leu2,3-112; lys2Δ; ura3-1; ADE2; trp1-1 | a | W303 | 1C;1D;2D;2E |
| CCY1744 | MATa/α; Bir1Δ/Δ::G418/HYG; ura3-1; LEU2/leu2,3-112; his3-11:pCUP1-GFP12-lacI12:HIS3 trp1-1:256lacO:TRP1 LYS2/lys2Δ; ADE2/ade2-1; can1-100; bar1Δ; pCC598::URA3 | a | W303 | S4A |
| CCY1905 | MATa; Bir1Δ:HYG; ura3-1; leu2,3-112: pCUP1-GFP12-lacI12:HIS3 trp1-1:256lacO: lys2Δ; can1-100 bar1Δ; pCC598::URA3 | b | W303 | 2B; 4B; 6A; S1C; S4C; S4D |
| CCY1934 | CCY1905 + UBP6::ubp6(E256X):G418 | c | W303 | S1C |
| CCY2999 | MATa Bir1Δ:HYG ; ura3-1; leu2;3-112: pCUP1-GFP12-lacI12:HIS3 trp1-1:256lacO:TRP1 lys2Δ ADE2 can1-100 bar1Δ | c | W303 | S4C |
| CCY3001 | CCY2999 + mps1-as1:G418 | c | W303 | S4C |
| CCY3031 | CCY1905 + duo1(P17L):G418 | c | W303 | 2B; 4B |
| CCY3033 | CCY1905 + cdc34(M64T):G418 | c | W303 | 2B |
| CCY3035 | CCY1905 + mif2(D241I):G418 | c | W303 | 2B |
| CCY3037 | CCY1905 + ndc80(K181N):G418 | c | W303 | 2B |
| CCY3039 | CCY1905 + spc105(R583G):G418 | c | W303 | 2B |
| CCY3041 | CCY1905 + sli15(G334S):G418 | c | W303 | 2B |
| CCY3043 | CCY1905 + sli15(L71S):G418 | c | W303 | 2B |
| CCY3047 | CCY1905 + cdc4(G439S):G418 | c | W303 | 2B; 6A |
| CCY3048 | CCY1905 + cdc4(S438G):G418 | c | W303 | 2B |
| CCY3050 | CCY1905 + mps1(R596H):G418 | c | W303 | 2B |
| CCY3052 | CCY1905 + mps1(V631M):G418 | c | W303 | 2B; S4C; S4D |
| CCY3064 | CCY1905 + spc34(D119A):G418 | c | W303 | 2B |
| CCY3083 | CCY1905 + rtg2(A433P):G418 | c | W303 | 2B |
| CCY3085 | CCY1905 + spc97(S816X):G418 | c | W303 | 2B |

| | | | | |
|---|---|---|---|---|
| CCY3210 | MATa/α; Bir1Δ/Δ:HYG ; ura3-1/ura3-1; leu2;3-112: pCUP1-GFP12-lacI12:HIS3/leu2;3-112 trp1-1:256lacO:TRP1/trp1-1:256lacO: lys2Δ ade2-1/ADE2 can1-100/can1-100 bar1Δ/bar1Δ; pCC598::URA3 DUO1/duo1(P17L):G418 | c | W303 | S4A |
| CCY3211 | MATa/α; Bir1Δ/Δ:HYG ; ura3-1/ura3-1; leu2;3-112: pCUP1-GFP12-lacI12:HIS3/leu2;3-112 trp1-1:256lacO:TRP1/trp1-1:256lacO: lys2Δ ade2-1/ADE2 can1-100/can1-100 bar1Δ/bar1Δ; pCC598::URA3 CDC4/cdc4(G439S):G418 | c | W303 | S4A |
| CCY3212 | MATa/α; Bir1Δ/Δ:HYG ; ura3-1/ura3-1; leu2;3-112: pCUP1-GFP12-lacI12:HIS3/leu2;3-112 trp1-1:256lacO:TRP1/trp1-1:256lacO: lys2Δ ade2-1/ADE2 can1-100/can1-100 bar1Δ/bar1Δ; pCC598::URA3 MPS1/mps1(V631M):G418 | c | W303 | S4A |
| CCY3299 | MATa; ura3-1; leu2;3-112; his3-11; trp1-1; ade2-1; lys2; ipl1-321; duo1(P17L):G418 | c | W303 | 3A |
| CCY3356 | MATa; ura3-1; leu2;3-112; his3-11; trp1-1; ade2-1; lys2; ipl1-321; cdc4(G439S):G418 | c | W303 | 3A |
| CCY3358 | MATa; ura3-1; leu2;3-112; his3-11; trp1-1; ade2-1; lys2; ipl1-321; mps1(V631M):G418 | c | W303 | 3A |
| CCY3439 | MATa/α; Bir1Δ/Δ:HYG ; ura3-1/ura3-1; leu2;3-112: pCUP1-GFP12-lacI12:HIS3/leu2;3-112 trp1-1:256lacO:TRP1/trp1-1:256lacO: lys2Δ ade2-1/ADE2 can1-100/can1-100 bar1Δ/bar1Δ; pCC598::URA3 DUO1/duo1Δ::NAT | c | W303 | S4A |
| CCY3441 | MATa/α; Bir1Δ/Δ:HYG ; ura3-1/ura3-1; leu2;3-112: pCUP1-GFP12-lacI12:HIS3/leu2;3-112 trp1-1:256lacO:TRP1/trp1-1:256lacO: lys2Δ ade2-1/ADE2 can1-100/can1-100 bar1Δ/bar1Δ; pCC598::URA3 CDC4/cdc4Δ::NAT | c | W303 | S4A |
| CCY3443 | MATa/α; Bir1Δ/Δ:HYG ; ura3-1/ura3-1; leu2;3-112: pCUP1-GFP12-lacI12:HIS3/leu2;3-112 trp1-1:256lacO:TRP1/trp1-1:256lacO: lys2Δ ade2-1/ADE2 can1-100/can1-100 bar1Δ/bar1Δ; pCC598::URA3 MPS1/mps1::NAT | c | W303 | S4A |
| CCY3445 | MATα; ura3-1; leu2;3-112; his3-11; trp1-1; ade2-1; ura3-1; leu2;3-112; his3-11; trp1-1; ade2-1; LYS2+; BUB3-mNeonGreen:NAT; NUF2-mRuby3:Hygro | c | W303 | 3B; 4C; S4B |
| CCY3469 | CCY1905 + dam1-765(S221F):G418 | c | W303 | S4D |
| CCY3643 | CCY3445 + duo1(P17L):G418 | c | W303 | 3B; 4C |
| CCY3644 | CCY3445 + cdc4(G439S):G418 | c | W303 | 3B |
| CCY3645 | CCY3445 + mps1(V631M):G418 | c | W303 | 3B; S4B |
| CCY3646 | CCY3445 + dam1-3D:G418 | c | W303 | 4C |
| CCY3684 | MATa ura3-1; leu2;3-112; his3-11; trp1-1; ade2-1; cdc4(G439S):G418 | c | W303 | S2A |
| CCY3688 | CCY1905 + dad1(N43S):G418 | c | W303 | 2B |
| CCY3694 | CCY1905 + ask1(S216F):G418 | c | W303 | 2B |
| CCY3730 | MATa; ura3-1; leu2;3-112; his3-11; trp1-1; ade2-1; LYS2+; mps1(V631M):G418 | c | W303 | S2A |
| CCY3736 | MATa; ura3-1; leu2;3-112; his3-11; trp1-1; ade2-1; LYS2+; dam1(3D):G418 | c | W303 | 4D |
| CCY3745 | MATa; ura3-1; leu2;3-112; his3-11; trp1-1; ade2-; LYS2+; duo1(P17L):G418 | c | W303 | 4D; S2A; S2C |
| CCY3750 | CCY1905 + dad2(K11Q):G418 | c | W303 | 2B |
| CCY3808 | MATα; ura3-1; leu2;3-112; his3-11; trp1-1; ade2-1; | c | W303 | 5B; 5C; |

| | | | | |
|---|---|---|---|---|
| | LYS2+; NUF2-mRuby3:Hygro; DAD3-mNeonGreen:NAT | | | S3 |
| CCY3809 | CCY3808 + duo1(P17L):G418 | c | W303 | 5B; 5C; S3 |
| CCY3813 | MATα; ura3-1; leu2;3-112; his3-11; trp1-1; ade2-1; LYS2+; NUF2-mRuby3:Hygro; DAD3-mNeon:NAT; CDC4::cdc4(G439S):G418 | c | W303 | 5B; 5C; S3 |
| CCY3816 | CCY3808 + mps1(V631M):G418 | c | W303 | 5B; 5C; S3 |
| CCY3821 | CCY3808 + dad1(N43S):G418 | c | W303 | 5B; 5C; S3 |
| CCY3880 | MATa; ura3-1; leu2;3-112; his3-11; trp1-1; ade2-1; lys2Δ; ipl1-321 | c | W303 | 3A |
| CCY3910 | CCY3445 + dam1(S20D):G418 | c | W303 | 4C |
| CCY3917 | CCY1905 + dam1(S218A;S221A):G418 | c | W303 | S4D |
| CCY3921 | MATa; ura3-1; his3-11, ade2-1; lys2Δ; can1-100 bar1Δ; ipl1-321; PDS1-18myc::LEU2; trp1-1:lacO:TRP1; dad2(K11Q):G418 | c | W303 | 3A |
| CCY3950 | MATa; ura3-1; leu2;3-112; trp1-1:256lacO: lys2Δ; ade2-1; can1-100 bar1Δ; pCUP1-GFP12-lacI12:HIS3; ipl1-321; dad1(N43S):G418; | c | W303 | 3A |
| CCY4093 | MATa, ura3-1; leu2;3-112; his3-11; trp1-1; dam1(S20D):G418 | c | W303 | 4D; S2C |
| CCY4106 | MATa; ipl1-321; PDS1-18myc::LEU2; trp1-1:lacO:TRP1; ura3-1; leu2;3-112: pCUP1-GFP12-lacI12:HIS3 trp1-1:256lacO: lys2Δ; spc34(D119A):G418 | c | W303 | 3A |
| CCY4108 | CCY3813 + dad2(K11Q):G418; | c | W303 | 5B; 5C; S3 |
| CCY4201 | MATα; ura3-1; leu2;3-112; his3-11; trp1-1; ade2-1; Sli15-6HA::NAT; Ipl1-3HA::HYG | c | W303 | S5 |
| CCY4254 | CCY3808 + dam1(S20D):G418 | c | W303 | 5B; 5C; S3 |
| CCY4271 | MATa PDS1-18myc::LEU2; ade2-1; his3-11; dam1(S20D):G418; DAD3-mNeon:NAT; duo1(P17L):G418 | c | W303 | S2C |
| CCY4329 | MATa; ura3-1; leu2;3-112; his3-11; trp1-1; ade2-1; Sli15-6HA::NAT; Ipl1-3HA::HYG; cdc4(G439S):G418; | c | W303 | S5 |
| CCY4834 | MATa; ura3-1; leu2;3-112; his3-11; trp1-1; ade2-1; dad1(N43S):G418 | c | W303 | S2A |
| CCY4854 | MATα; ura3-1; leu2;3-112; his3-11; trp1-1; ade2-1; BUB3-mNeonGreen:NAT; NUF2-mRuby3:Hygro; dad1(N43S):G418 | c | W303 | 3B |
| CCY4863 | MATα; ura3-1; leu2;3-112; his3-11; trp1-1; ade2-1; BUB3-mNeonGreen:NAT; NUF2-mRuby3:Hygro; spc34(D119A):G418 | c | W303 | 3B |
| CCY4911 | MATa; ura3-1; leu2;3-112; his3-11; trp1-1; ade2-1; pRS316::URA3 | c | W303 | 2C; 3C; 4E; S2D |
| CCY4913 | CCY4911 + duo1(P17L):G418 | c | W303 | 3C; 4E; S2D |
| CCY4915 | CCY4911 + dad1(N43S):G418 | c | W303 | 3C |
| CCY4921 | CCY4911 + cdc4(G439S):G418 | c | W303 | 3C |
| CCY4923 | CCY4911 + mps1(V631M):G418 | c | W303 | 3C |
| CCY4925 | CCY4911 + dam1(S20D):G418 | c | W303 | 4E; S2D |
| CCY4927 | CCY4911 + dam1(3D):G418 | c | W303 | 4E |
| CCY4929 | MATa; PDS1-18myc::LEU2; | c | W303 | S2D |

| | | | | |
|---|---|---|---|---|
| | DAM1::dam1(S20D):G418; DAD3-mNeonGreen:NAT; DUO1::duo1(P17L):G418; pRS316::URA3 | | | |
| CCY4933 | CCY4968 + duo1(P17L):G418 | c | W303 | 2C |
| CCY4935 | CCY4968 + dad1(N43S):G418 | c | W303 | 2C |
| CCY4937 | CCY4968 + dad2(K11Q):G418 | c | W303 | 2C |
| CCY4939 | CCY4968 + spc34(D119A):G418 | c | W303 | 2C |
| CCY4947 | CCY4968 + mps1(V631M):G418 | c | W303 | 2C |
| CCY4949 | CCY4968 + cdc4(G439S):G418 | c | W303 | 2C |
| CCY4968 | MATa; Bir1Δ:HYG; ura3-1; leu2;3-112: pCUP1-GFP12-lacI12:HIS3 trp1-1:256lacO: lys2Δ; can1-100 bar1Δ; pRS316::URA3 | c | W303 | 2C |
| CCY4997 | MATa; Bir1Δ:HYG; ura3-1; leu2;3-112: pCUP1-GFP12-lacI12:HIS3; can1-100; bar1Δ; pCC598::URA3; leu2;3-112; trp1-1:lacO:TRP1; lys2Δ; dam1(S20D):G418 | c | W303 | 4B |
| CCY5001 | MATa; Bir1Δ:HYG; ura3-1; leu2;3-112: pCUP1-GFP12-lacI12:HIS3; trp1-1; pCC598::URA3; dam1(3D):G418 | c | W303 | 4B |
| CCY5101 | MATa/α; ura3-1; leu2;3-112; his3-11; trp1-1; ade2-1; ura3-1; leu2;3-112; his3-11; trp1-1; DAM1/dam1(S20D):G418; DUO1/duo1(P17L):NAT | c | W303 | S2B |
| CCY5107 | MATa/α; ura3-1; leu2;3-112; his3-11; trp1-1; ade2-1; ura3-1; leu2;3-112; his3-11; trp1-1; DAM1/dam1(S20D):G418; CDC4/cdc4(G439S):NAT | c | W303 | S2B |
| CCY5109 | MATa/α; ura3-1; leu2;3-112; his3-11; trp1-1; ade2-1; ura3-1; leu2;3-112; his3-11; trp1-1; DAM1/dam1(S20D):G418; MPS1/mps1(V631M):NAT | c | W303 | S2B |
| CCY5119 | MATa/α; ura3-1; leu2;3-112; his3-11; trp1-1; ade2-1; ura3-1; leu2;3-112; his3-11; trp1-1; DAM1/dam1(S20D):G418; DAD1/dad1(N43S):NAT | c | W303 | S2B |
| CCY5282 | MATa; ura3-1; leu2;3-112; his3-11; trp1-1; ade2-1; pRS316::URA3; ctf19Δ:G418 | c | W303 | 2C; 3C; 4E; S2D |
| CCY5326 | MATa; ura3-1; leu2;3-112; his3-11; trp1-1; ade2-1; ADE2; Nuf2-mCherry::G418; pGal1-3HA-Sli15::HIS3; mNeonGreen-6HA-Sli15::URA3; pGal-Sgo1::HYG | c | W303 | 6B |
| CCY5373 | CCY3445 + dad2(K11Q):G418 | c | W303 | 3B |
| CCY5376 | MATa; ura3-1; leu2;3-112; his3-11; trp1-1; ade2-1; spc34(D119A):G418 | c | W303 | S2A |
| CCY5377 | MATa; ura3-1; leu2;3-112; his3-11; trp1-1; ade2-1; dad2(K11Q):G418 | c | W303 | S2A |
| CCY5420 | MATa; ura3-1; leu2;3-112; his3-11; trp1-1; ade2-1; spc34(D119A):G418; pRS316::URA3 | c | W303 | 3C |
| CCY5422 | MATa; ura3-1; leu2;3-112; his3-11; trp1-1; ade2-1; dad2(K11Q):G418; pRS316::URA3 | c | W303 | 3C |
| CCY5424 | MATa/α; Bir1Δ/Bir1Δ:HYG; ura3-1; leu2,3-112: pCUP1-GFP12-lacI12:HIS3 trp1-1:256lacO: lys2Δ can1-100 bar1Δ; pCC598::URA3 | c | W303 | S4A |
| CCY5435 | MATa; ura3-1; leu2;3-112; his3-11; trp1-1; ade2-1; ADE2; Nuf2-mCherry::G418; pGal1-3HA-Sli15::HIS3; mNeonGreen-6HA-Sli15::URA3 | c | W303 | 6B |
| CCY5469 | MATa; ura3-1; leu2;3-112; his3-11; trp1-1; ade2-1; ADE2; Nuf2-mCherry::G418; pGal1-3HA-Sli15::HIS3; mNeonGreen-6HA-Sli15::URA3; pGal-Sgo1::HYG cdc4(G439S):NAT | c | W303 | 6B |
| CCY5508 | MATa; ura3-1; leu2;3-112; his3-11; trp1-1; ade2-1; ADE2; Nuf2-mCherry::G418; pGal1-3HA-Sli15::HIS3; | c | W303 | 6B |

| | | | | |
|---|---|---|---|---|
| | mNeonGreen-6HA-Sli15::URA3; cdc4(G439S):NAT | | | |
| CCY5512 | MATa/α; ura3-1; leu2;3-112; his3-11; trp1-1; ade2-1; ura3-1; leu2;3-112; his3-11; trp1-1; dad2(K11Q):NAT; dam1(S20D):G418 | c | W303 | S2B |
| CCY5518 | MATa/α; ura3-1; leu2;3-112; his3-11; trp1-1; ade2-1; ura3-1; leu2;3-112; his3-11; trp1-1; spc34(D119A):NAT; dam1(S20D):G418 | c | W303 | S2B |
| CCY5524 | CCY3808 + spc34(D119A):G418 | c | W303 | 5B; 5C; S3 |
| CCY5532 | CCY1905 + met30-6:G418 | c | W303 | 6A |
| CCY5535 | CCY1905 + cdc4-1:G418 | c | W303 | 6A |

| **Plasmid** | **Description** | **Source** |
|---|---|---|
| pRS306 | pBluescript + URA3 | d |
| pRS316 | pBluescript + URA3 | d |
| pCC598 | BIR1 + 1000 bases upstream in pRS306 | b |
| pCC-H046 | Repair plasmid for $FBXW7^{G437S}$ | c |

| **Cell line** | **Description** | **Source** |
|---|---|---|
| Clone C3 | HAP1 $TP53^{-}$ | e |
| CCH2248 | HAP1 $TP53^{-}$, $FBXW7^{G437S}$ (clone 1) | c |
| CCH2251 | HAP1 $TP53^{-}$, $FBXW7^{G437S}$ (clone 2) | c |
| CCH2254 | HAP1 $TP53^{-}$, $FBXW7^{G437S}$ (clone 3) | c |

a – Campbell and Desai; Nature 497: 118-121 (2013)
b – Ravichandran et al; Genes and Development (2018)
c – This study
d – Sinorski and Hieter Genetics 122: 19-27 (1989)
e – J. Loizou lab
